# Protein target search diffusion-association/dissociation free energy landscape around DNA binding site with flanking sequences

**DOI:** 10.1101/2024.08.26.609820

**Authors:** Biao Wan, Jin Yu

## Abstract

Proteins including transcription factors (TFs) and regulating enzymes search DNA for specific sites by alternating 3D diffusion in cell nucleus space and 1D diffusion on DNA. The search dynamics and free energy landscape of the protein along DNA depend essentially on the protein-DNA interactions, which simultaneously determine the protein-DNA association strength and relative population profiling along DNA, e.g., measured from protein binding microarray (PBM) to genome-wide mapping. Here we present a minimal structure-based model of protein diffusional search along DNA amid protein binding and unbinding events on the DNA, taking into account protein-DNA electrostatic interactions and hydrogen-bonding (HB) interactions or contacts at the interface. We accordingly constructed the protein diffusion-association/dissociation free energy surface and mapped it to 1D as the protein slides along DNA, maintaining the protein-DNA interfacial HB contacts that presumably dictate the DNA sequence information detection. Upon DNA helical path correction, the protein 1D diffusion rates along DNA can be physically derived to be consistent with experimental measurements. We also show that the sequence-dependent protein sliding or stepping patterns along DNA are regulated by collective interfacial HB dynamics, which also determines the ruggedness of the 1D diffusion free energy landscape. In comparison, protein association or binding with DNA are generically dictated by the protein-DNA electrostatic interactions, with an interaction zone of nanometers around DNA. Extra degrees of freedom (DOFs) of the protein such as rotations and conformational fluctuations can be well accommodated within the electrostatic interaction zone. As such we demonstrate that the protein binding or association free energy profiling along DNA smoothens over the 1D diffusion free energy landscape, which leads to population variations for an order of magnitude upon a marginal free energetic smoothening around the specific or consensus sites. We further show that the protein unbinding or dissociation from a comparatively high-binding affinity DNA site is dominated by lateral diffusion to the flanking low-affinity sites. The results suggest that experimental characterizations on the relative protein-DNA binding affinities or population profiling on the DNA are systematically and physically impacted by the extra DOFs of protein motions aside from translation as well as from flanking DNA sequences due to protein 1D diffusion and non-specific binding/unbinding.

## I. INTRODUCTION

Protein-DNA interactions play essential roles in genetic and epigenetic regulations. Deciphering how proteins such as transcription factors (TFs) search efficiently and locate accurately the target sites are of high significances for both physical understanding and biomedical applications (1-5). Although the facilitated diffusion models laid foundation on quantitative characterizations of the TF-DNA target search, i.e., by suggesting alternating between 3D diffusion in the nucleus space and 1D diffusion on the DNA (6-10), the corresponding protein-DNA structural dynamics remain vague. While challenges on elucidating the *in vivo* search are due to highly crowded, heterogeneous, and fluctuating cellular environment in general, genome folding or packing in high organisms and miscellaneous protein-protein interactions and regulations add notable complications. Despite of high-resolution structural characterizations of regulatory protein-DNA complexes, single-molecule measurements on TF diffusional dynamics on DNA remain challenging for simultaneous sub-nanometer and sub-millisecond detections (11-15). On the other hand, structure-based molecular dynamics (MD) simulations started demonstrating protein-DNA dynamics in search and recognition from atomic to coarse-grained resolution (16 - 22). Nevertheless, the studies rely on high-performance computing (HPC) and are resource demanding, hardly approaching to or beyond millisecond long time scale. In current work, in order to present a theoretical framework computationally non-intensive while taking into account the most essential physical interactions in the protein-DNA target search, comparable between specific and non-specific binding sites (23), we employed a minimum structure-based model of a spherical protein and a linear DNA, locally, and constructed a free energy landscape of the protein diffusion-association/dissociation along DNA.

The free energy landscape is commonly employed to project the complex system dynamics in equilibrium ensemble from high-dimension to low-dimension (24-29). In current case, the free energy landscape is 2D, considering both directions of protein 1D diffusion (along DNA) and dissociation/association pathway (perpendicular to DNA), or 1D if considering only the protein 1D diffusion or the relative protein association profiling along DNA. Comparable to protein-folding free energy landscape, ruggedness of the landscape has been addressed in modeling works on the protein-DNA target search (5, 30-32). However, previous models were phenomenological by introducing the parameter of the ruggedness directly, instead of deriving it physically from modeling the protein-DNA interactions. In current work, we modeled the protein-DNA interactions explicitly, including the generic electrostatic interactions and sequence-dependent hydrogen bond (HB) interactions at the protein-DNA interface, both of which are present between protein and DNA no matter on specific or non-specific DNA sequences (17). By further considering the solution hydrodynamics with a DNA helical path correction (33), we derived the proper time scales of the protein 1D diffusion along DNA, consistently with experimental measurements (33). Consequently, the local ruggedness of the protein 1D diffusion free energy landscape well reflect the sequence-dependent HB interactions modeled at the protein-DNA interface, aside from the generic protein-DNA electrostatic interactions, which depend the effective protein charges, geometrical features of protein-DNA, and essentially the solution ionic conditions.

In current model, besides analytically and numerically characterizing the protein diffusional search dynamics on the DNA, from sub-nanosecond to sub-millisecond time scale, protein unbinding or dissociation events from DNA around millisecond time scale are also accounted and sampled. In particular, current studies relate the protein 1D diffusion and binding/unbinding dynamics to the protein-DNA interfacial HB and electrostatic interactions, respectively. One can thus compare the protein 1D diffusion landscape, which contains most abundant DNA sequence information as protein closely diffuses or slides along DNA, with the protein relative binding or association profiling along DNA, which can be measured *in vitro* by protein binding microarray (PBM) (34, 35) or *in vivo* from genome-wide mapping such as ChIP-seq (36, 37). By illustrating the protein 1D diffusion free energy landscape dictated by collective HB dynamics at the protein-DNA interface, we predict sequence-dependent protein stepping characteristics along DNA, which can be tested experimentally. We also relate various protein stepping sizes (>1-bp) along random DNA with comparatively high binding-affinities DNA consensus sites, which can also subject to future experimental tests. Furthermore, we attribute the dominant path of protein dissociation from DNA to the protein lateral diffusion to the flanking DNA sequence, so that to explain the flanking sequence impacts (38-40), i.e., via a physical perspective.

## II. METHOD AND MODEL CONSTRUCTION

### A. Physical interactions modeled between protein and DNA

The TF domain protein is modeled as a charged sphere with rotational DOFs (**FIG 1A**). The electrostatic charge distribution and polarization of the protein can be represented by placing effectively the positive and negative charges on the surface of the spherical protein. To keep the protein electrostatically neutral, approximately, one can place an equal number of positive and negative charges on the lower and upper hemispheres, respectively, with the lower positively charged hemisphere bound to the negatively charged linear DNA. On the other hand, protein binding sites on DNA are discretized and linearly arranged with a spacing distance of one base pair (bp) at ∼ 0.34 nm (**FIG 1B**). The protein-DNA electrostatic interactions with solution screening by ions can be approximated by the Debye-Huckel potential

**FIG 1.**
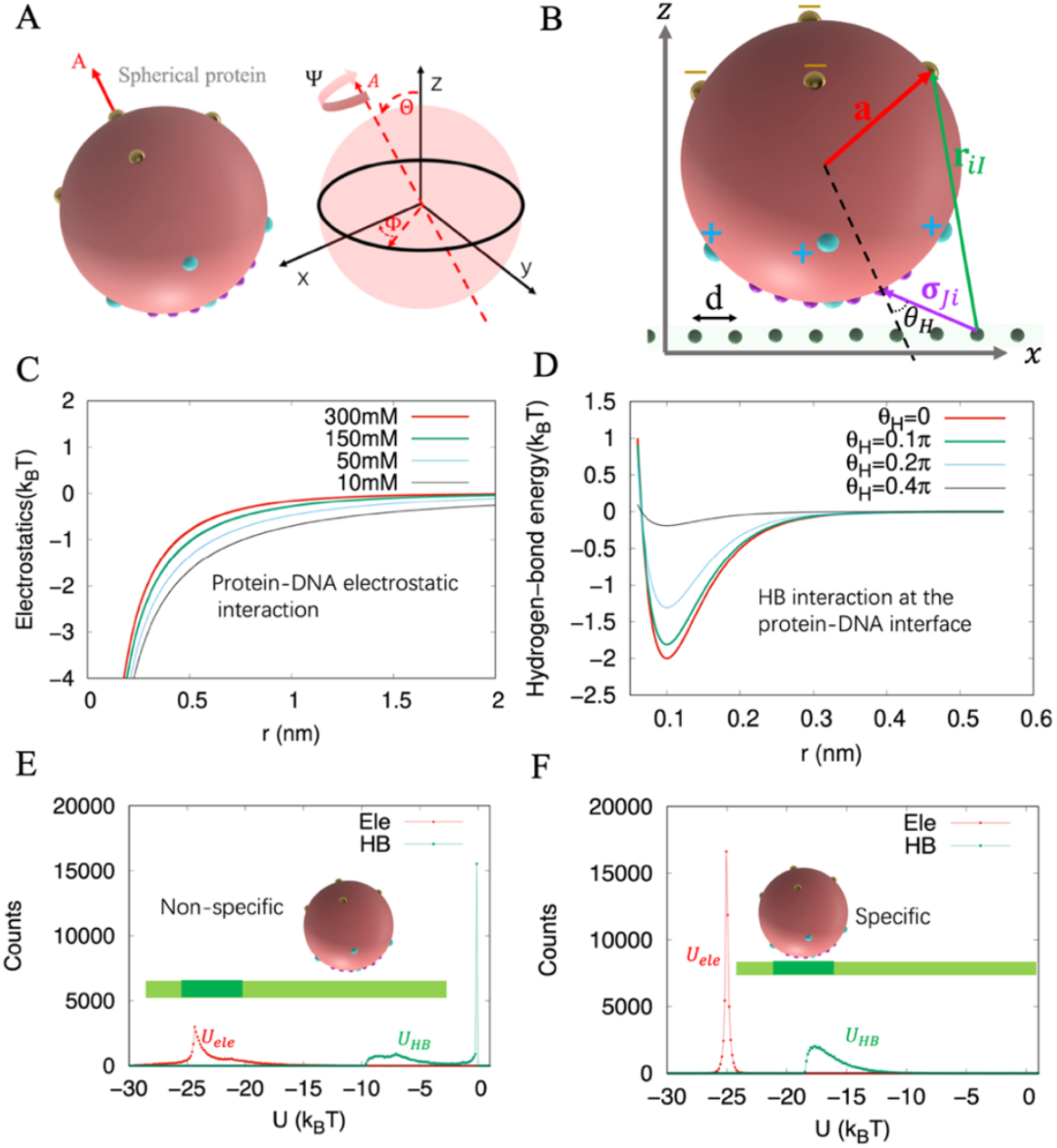
Model of a spherical protein interacting with a linear DNA. (A) Schematic of the rotational DOFs (Θ, Φ, Ψ) of the spherical protein. (B)The effective positive and negative charges are indicated by cyan and coral dots, respectively. The HB interaction sites (violet dots) are mounted on the positively charged hemisphere. Binding charges on DNA are embedded in a tube with a diameter of 0.2 *nm* and aligned on the x-axis separated individually by *d* = 0.34 *nm* in distance. ***r***_*iI*_ stands for the vectors between a DNA binding site and an interaction site on the protein, and ***a*** is the radial vector in spherical coordinates. HB angle *θ*_*H*_ is measured between the radial direction (dashed line) of the HB interaction site on the protein sphere and the direction (solid line) of the pair-interaction, i.e., *cosθ*_*H*_ = −***a*** · ***σ***_*iJ*_. (C) Protein-DNA electrostatic interactions modeled via the Debye-Huckel potential under different solution ionic strengths. (D) HB interaction modeled via the Morse potential with orientation dependence on bond angle *θ*_*H*_, and with strength ℰ_*HB,iJ*_ sequence encoded by PWM. (E)&(F) Electrostatic and HB energies of protein-DNA interactions on non-specific and specific DNA binding sites.

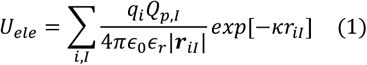

where *q*_*i*_ = −2*e*_*i*_ is the efective charge of one pair of nucleotides on the *i*-th site of DNA, *Q*_*p,I*_ is the efective charge of the *I*-th charged site on the spherical protein, *ϵ*_*r*_ = 78.25 is the dielectric constant of water (41), ***r***_*iI*_ the radial vectors between the *i*-th DNA binding site and the *I*-th charged site on the protein (**FIG 1B**), and *κ* the inverse Debye length, as a function of ionic strength *I*_s_, i.e., 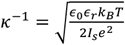 characterizing the strength of the electrostatic screening (**FIG 1C**) (42). For example, *k*^−1^∼ 0.8 *nm* for *I*_s_ ∼150 mM ionic strength in physiological condition. With minor variations of the ionic strengths, the electrostatic interactions converge from ∼ 1-2 nm away from DNA (**FIG 1C**).

Aside from the electrostatics, the HB interaction sites are placed on the positively charged protein hemisphere and the linear DNA, and the HBs are to be formed at the protein-DNA interface. We employ the *Morse* potential accompanied with the bond angle to account for the HB interaction empirically(**FIG 1D**) (43, 44)

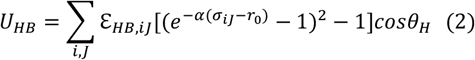

where ℰ_*HB,J*_ is the depth of the Morse potential of the *J*-th HB site on protein, *α* ∼ 10^−1^*nm*, is the inverse decay length, *r*_0_ ∼ 10^−1^*nm*, specifies the radius of the repulsive core, *σ*_*iJ*_ the distance between the *i*-th DNA binding site and the *J*-th HB site on the protein (**FIG 1B**), and *θ*_*H*_ is the bond angle. Within a distance ∼0.4 nm, the HB interaction is sensitive to the bond angle or direction (**FIG 1D**). In this study, the strength ℰ_*HB,iJ*_ is set sequence-dependent, where *i* takes 1 to 4, labeling for ATCG, and *J* from 1 to *n*, denoting the protein-DNA HB interaction sitesc, with *n=6* used by default below. One can define the strength of HB interaction according to a position weight matrix (PWM) (45, 46), i.e., obtained from large-scale sequence analyses of protein binding preferences. For illustration purpose, we use the PMW shown below, e.g., to characteriz a W-box sequence ‘**TTGACT**’ for a small TF domain protein WRKY binding (17, 47),

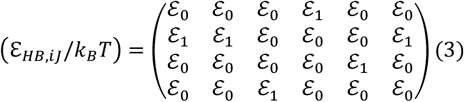

where we used ℰ_0_ = 0.8 and ℰ_1_ = 7.2 (with energy units *k*_*B*_*T*) for non-preferred and preferred HB contacts, according to the PWM, between the protein and various DNA sequences.

Accordingly, we take into the electrostatics and HB interactions between protein and DNA into the total energy function as,

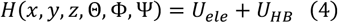

where the coordinates x, y, and z describe the location of the protein center of mass (COM) on the DNA, which is modeled linearly along x; Θ, Φ and Ψ denote the protein orientations, with {Θ, Φ} the spatial angle of the protein ‘polar’ axis in spherical coordinates, and Ψ the rotation or ‘spin’ of the protein around its own polar axis. It should be noted that the protein COM is confined in the z-x plane under an assumption of rotational symmetry of DNA around its long axis or the x-axis, and DNA is modeled linear locally (15 to 50 bp) as it is much shorter than the persistence length (∼150 bp). In current model, we focus on protein-DNA interactions in the presence of protein rotational motions, while ignoring protein conformational changes and DNA coiling dynamics on large scale. Though the electrostatics dominate the overall protein-DNA interactions, one sees that the HB interactions at protein-DNA interface essentially distinguish the specific and non-specific DNA binding sites (**FIG 1E&F**).

### B. Langevin dynamics of the spherical protein

According to the total energy function from Eq (4), a dynamic model of the spherical protein moving along DNA is developed. To allow sufficient samplings, the length of DNA used by default is 15-bp, and the DNA is kept immobile. Consequently, the COM of the protein moves along DNA following the Langevin equation dictated by the protein-DNA interaction,

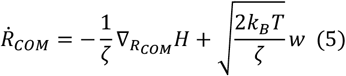

where *ζ* = 6*πηa* is the translational friction coefficient, with *η* the solution viscosity, and *a* the radius of the spherical protein; *k*_*B*_ is the Boltzmann constant, and *w*(t) is the Gaussian noise with zero mean and unit variance. The orientational or rotational motions of the protein due to the protein-DNA interactions and thermal fluctuations are modeled by separating the spatial rotation of the unit axial vector **A** (i.e., the protein polar axis), Δ**Ω**_*A*_ on {Θ, Φ} from the protein spin about **A**, Ψ. Accordingly, the protein orientational Langevin dynamic equations are formulated as (details in **Supplementary Material S1**):

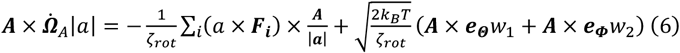

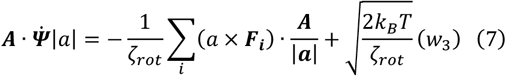

where *ζ*_*rot*_ = 8*πηa* is the friction coefficient for the rotation of the protein in solution, and *w*_1_, *w*_2_ and *w*_3_ are the Gaussian noise with zero mean and unit variance.

## III. RESULTS

### A. Construction of protein diffusion-association/dissociation free energy landscape

To characterize both protein 1D diffusion along DNA (*x* direction) and occasional dissociation/unbinding from DNA (*z* direction), we constructed the potential of mean forces (PMF) in 2D coordinate space as *U*(*x, z*), which provides a unified free energy surface describing both the diffusion and binding-unbinding dynamics of protein. Projection of the 2D potential to the 1D diffusion (*x*) or mapping it along the unbinding (*δz* >0) direction, leads to the diffusional free energy landscape Δ*G*_*diff*_(*x, z*_0_) on the DNA (*δz* ∼ 0 or at *z*∼*z*_0_), or the dissociation free energy profile *φ*(*x*′, *δz* > 0) at *x* = *x*′ and the binding free energetics between the *bound* and the *unbound* states as ΔG_*b*_ (x′) ≅ *φ*(*δz* < *δz*_*E*_) − *φ*(*δz* > *δz*_*E*_), where *δz*_*E*_ denotes the protein-DNA *electrostatic interaction zone* (about 2-3 nm). Further, one obtains the *relative* protein binding profiling along DNA as ΔΔ*G*_*b*_(*x*) according to ΔG_*b*_(x). As a result, Δ*G*_*diff*_(*x*) describes the protein 1D diffusion along DNA as the protein-DNA interfacial HB interactions are well maintained (*z*∼*z*_0_), while ΔΔ*G*_*b*_(*x*) reflects the protein association profiling or changes of the binding free energetics along DNA due to the protein-DNA electrostatic interactions, *with* or *without* the interfacial HBs. Besides, the protein rotation DOFs also affect significantly the protein-DNA interaction energetics. Below we first constructed the *U*(*x, z*), ΔΔ*G*_*b*_(*x*), and Δ*G*_*diff*_(*x*) analytically, assuming protein rotational DOFs either very fast (integrated out) or very slow (with fixed orientation). Later, we employed the Langevin dynamics with both translation and rotation DOFs of the protein along DNA, and provided numerical solutions.

#### 1. Protein-DNA interaction PMF or 2D potential surface U(x, z)

Assuming that the protein rotational DOFs are independent of the translation motions, the mean force to resist the protein dissociation from DNA along the positive *z*-axis is written as

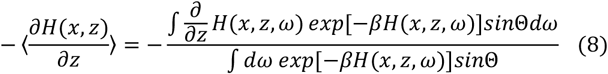

where *ω* ≡ {Θ, Φ, Ψ} denotes the rotation DOFs and *dω* = *d*Θ*d*Φ*d*Ψ. During protein translational search along or away from DNA, the rotational DOFs are not necessarily following or relaxed at each (x,z) position, and we address the issues below.

If the rotational motions of the protein are fast, i.e., to be completely relaxed at any spatial position (x, z), one integrates fully over 0 ≤ Θ < *π*, 0 ≤ Φ < *π* and 0 ≤ Ψ < 2*π* in Eq (8). In such a case, the 2D molecular interaction potential or PMF at a given position *(x, z*), i.e., for protein diffusing on the DNA (*z* = *z*_0_) or dissociating from DNA (to *z* = ∞ or sufficient large) is defined as:

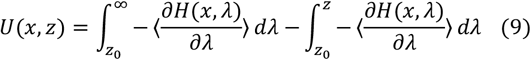

where *z*_0_ (>0) denotes the location of the protein COM when the protein is bound to the DNA, or due to the protein size; *λ* stands for the location parameter along the *z*-axis. The molecular interaction potential *U*(*x, z*) thus represents the energetic projection that fully averages out the fast-angular motions of (Θ, Φ, Ψ). With that, one can describe the protein dissociation and diffusion dynamics surrounding and along DNA. The potentials are visualized in **FIG 2**.

**FIG. 2:**
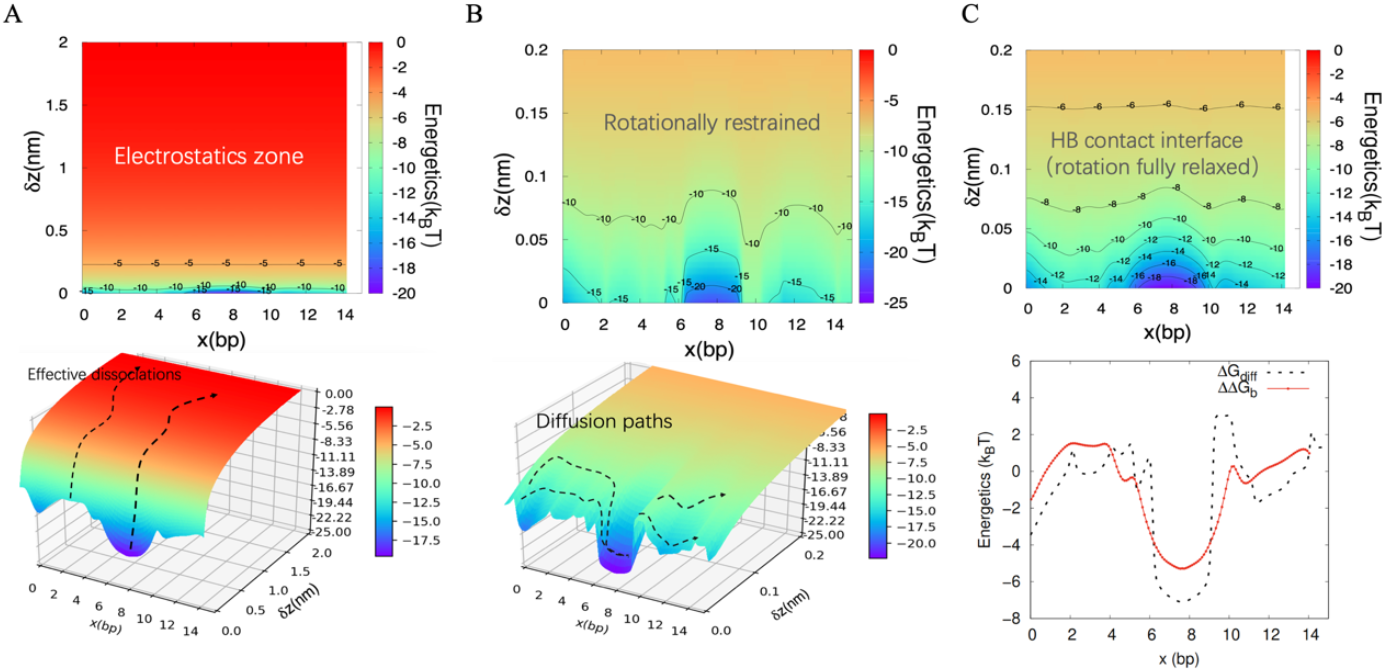
Construction of the protein diffusion-association/dissociation free energy landscape according to the protein-DNA interaction potential for protein diffusion and association along DNA. (A) *U*(*x, δz*) as a function of protein diffusion along DNA or *x*, and protein dissociation away from DNA *δz* ≡ *z* – *z*_0_ > 0 (*z*_0_ as the protein bound position on DNA). Color bars represent the depths of the potential (in *k*_*b*_*T*). The 15-bp DNA sequence is designated ‘GTCGT**<u>TTGACT</u>**TAAT’, with ‘**<u>TTGACT</u>**’ the specific protein-DNA binding site in the center, specified according to the PWM. Dissociations to infinity (*δz >* 2*nm*) can be effectively depicted as escaping from the electrostatic zone (*bottom*). (B) The potential *U*(*x, δz*) corresponding to protein diffusion on the DNA as maintaining the protein-DNA HB contacts (*δz <* 0.2*nm*). The protein rotational DOFs are partially restrained: The angular confinements for Θ is set at 0.1*π*, Ψ is −0.1*π* to 0.1*π* and Φ is free. The protein diffusion paths can be seen on the potential profile (*bottom*.) (C) The potential *U*(*x, δz*) on the HB contact surface (as in B) but free of angular restraints (rotation DOFs fully relaxed). A comparison of the 1D free energy landscape for protein diffusion Δ*G*_*diff*_ (as from B) and for the relative protein binding or association profiling ΔΔ*G*_*b*_ is also shown (*bottom*).

#### 2. Protein relative binding/association free energy profiling along DNA within the protein-DNA electrostatic interaction zone

Based on the 2D potential *U*(*x, z*), the protein binding free energetics (at position *x* along DNA) can be determined by calculating the energetics of the protein dissociation out of the electrostatic zone with DNA, vertically (along *z*-axis), and approaching to an effective infinity as the free energetics does not change further, or converges (*δz* ≡ *z* − *z*_0_ > *δz*_*E*_∼2 *nm*) (**FIG 2A**). Accordingly, the binding free energetics of protein bound at position *x* on the DNA can be measured via calculating the free energy change upon quasi-statistically pulling the protein along *z* from the initially bound position within the electrostatic zone (0< *z* − *z*_0_ < *δz*_*E*_) to the infinity (or effectively for *z* − *z*_0_ > *δz*_*E*_). The relative binding free energy profile of the protein along DNA, within the electrostatic interaction or association zone *δz*_*E*_, can be then written as:

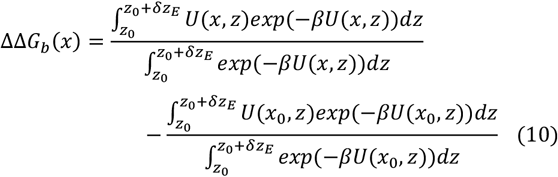

#### 3. Protein 1D diffusion free energy landscape at the protein-DNA HB contact interface

Protein diffusion or sliding on the DNA is defined as that the protein-DNA interface maintains abundant HB contacts (*δz* < *δz*_*H*_ < 1*nm*). In addition to the distance, the interfacial HB contacts can be easily broken if the protein undergoes rotation, though the protein remains in association with DNA, i.e., within the electrostatic zone. That is to say, in order to maintain HB contacts with DNA, the protein must undergo restrained orientations during diffusion along DNA. Based on numerical simulations, the integral upper bound of Θ takes 0.1*π*, the lower and upper bound of Ψ take −0.1*π* and 0.1*π* (see **S2 FIG S5**), and the HB broken or the cutoff distance is set as *δz*_*H*_ = 0.2 *nm*. To analytically show the diffusion free energy profile in **FIG 2B**, we first assume that the protein rotational degrees are partially restrained within certain bounds. As a result, the mean force for the protein to overcome the energy barriers for diffusion or sliding on the DNA, without breaking the HB contacts with DNA, is

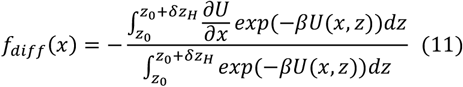

with *δ z*_*H*_ the cut-off distance for maintaining the HBs between the DNA and protein. Accordingly, the 1D diffusion free energy landscape along the *x*-axis is

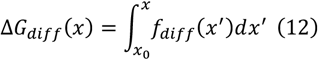

The corresponding potential is shown in **FIG 2B** for the protein rotation partially restrained case. The diffusion can thus be depicted as a walker wandering on the potential surface *U*(*x, z*) while maintaining the protein-DNA HB contacts (*δz* < *δz*_*H*_).

In comparison with the relative protein-DNA binding free energy profile ΔΔ*G*_*b*_, the protein-DNA diffusional free energy landscape is more rugged, e.g., with the energy minimum of Δ*G*_*diff*_ ∼ 2*k*_*B*_*T* lower than that of ΔΔ*G*_*b*_, and a maximal barrier ∼ 5 *k*_*B*_*T* higher than that of ΔΔ*G*_*b*_ (**FIG 2C**). It happens as the protein undergoes restricted angular displacements during diffusion to maintain the protein-DNA interfacial HB contacts, which would not be the case once the protein undergoes slight dissociation from DNA, i.e., breaking the HB contacts but remaining association with DNA or inside the protein-DNA electrostatic zone (*δz*_*H*_ < *δz* < *δz*_*E*_). Consequently, the protein-DNA binding profiling ΔΔ*G*_*J*_ smoothens over the 1D protein-DNA diffusion free energy landscape Δ*G*_*diff*_, as shown in **FIG 2C** (*bottom*).

### B. Langevin dynamics on protein diffusion along DNA and dissociation from DNA

We demonstrate first that by modeling protein-DNA electrostatic and HB interactions at the interface, one can well reproduce experimentally measured diffusion coefficients, upon DNA helical path corrections. By properly capturing the protein 1D diffusion time scale along DNA via the Langevin dynamics, we then show the sequence-dependent stepping statistics of the protein and collective interfacial HB dynamics during the diffusion, and further investigate stochastic events of the protein dissociation from the specific or consensus DNA binding sites. We demonstrate that the dissociation is often proceeded by the protein lateral diffusion to the flanking DNA sites.

#### 1. Protein-DNA physical interactions along with the helical path correction contribute to proper time scale of protein 1D diffusion ∼ bp^2^/ μs on the DNA

From results above, one sees that the protein-DNA electrostatic and HB interactions together contribute to the free energy barriers along DNA. The diffusional free energy landscape Δ*G*_*diff*_ describes the sliding of protein along DNA while maintaining abundant HBs at the protein-DNA interface, which essentially determine the effective diffusion coefficient

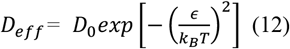

where *ϵ* is the *ruggedness* of the diffusion free energy profile and *D*_0_ is the 1D diffusion coefficient for protein in solvent conditions (48). Note that *ϵ* is commonly introduced as a phenomenological parameter, while below we show *ϵ* numerically as a consequence of the physical interactions between the protein and DNA.

For protein sliding along the helical strand or groove of DNA, aside from the protein-DNA interactions modeled above, an essential contribution to protein-DNA ‘friction’ comes from the protein curvilinear motion along the DNA helical path, along with the protein spinning about its own axis (**FIG 3A**). Taking into account these effects, the overall friction coefficient is given by (49, 50)

**FIG. 3:**
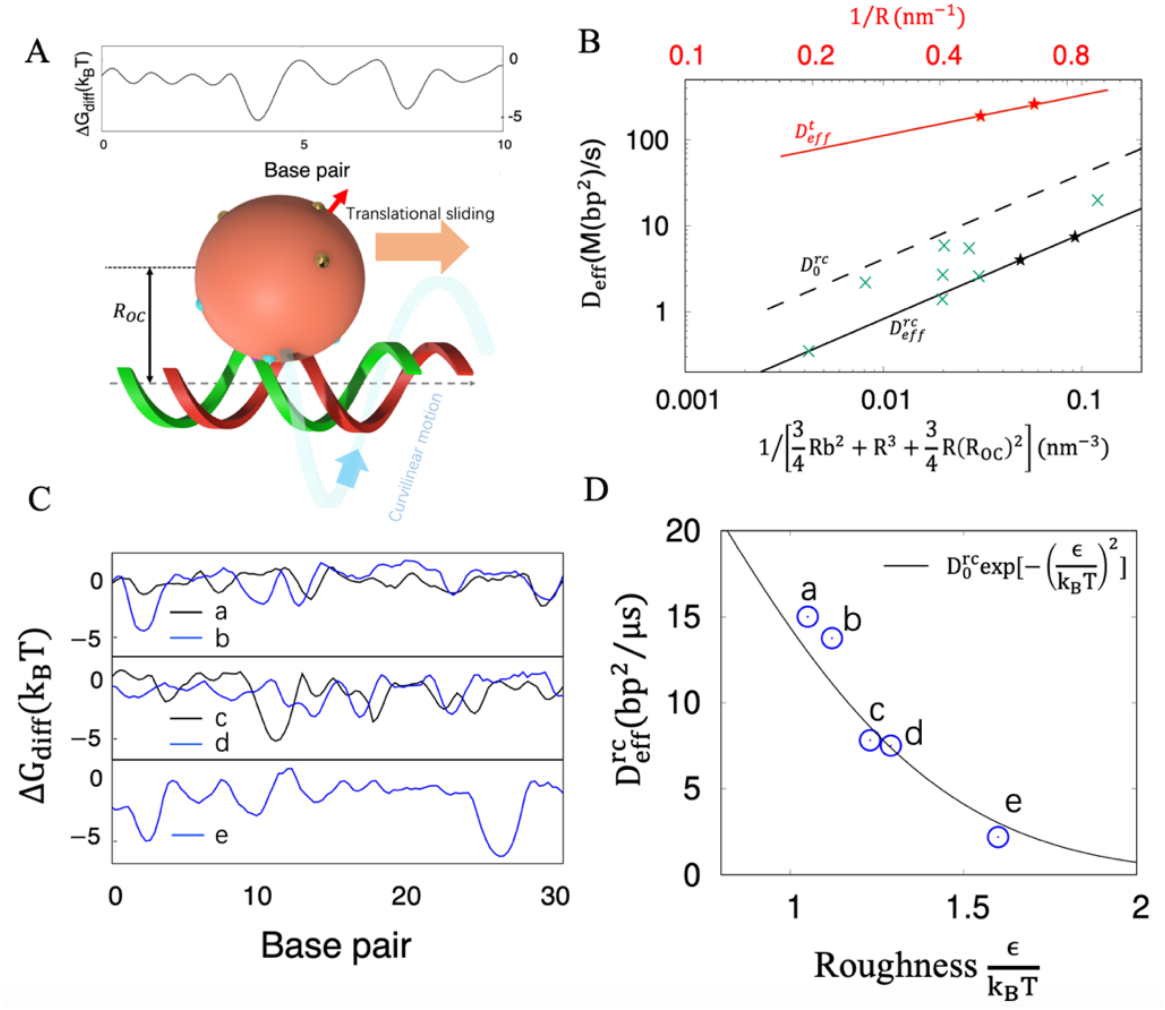
Deriving the protein 1D effective diffusion rates along DNA. (A) Schematics of protein 1D diffusion free energy landscape and the spherical protein on the DNA helical path. (B) Effective diffusion coefficients of the spherical protein for pure translational sliding 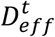 (red; without helical-path correction) and for rotation-coupled sliding (black; with helical-path correction: 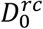 (dashed line) and 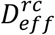 (solid line) are the 1D diffusion coefficient for the protein rotation-coupled sliding in regular solvent condition (no protein-DNA interaction) and along DNA (with the protein-DNA interactions), respectively. The derived effective diffusion coefficients for *R* = 1.5*nm* and *R* = 2*nm* are labeled as stars. The diffusion coefficients for different proteins from the experimental study are shown as the green points (49). (C) The local diffusion free energy landscape for 5 random DNA sequences (30-bp for each; a:TTAGCTTATCCCACCAAGCTTCAGCATGCAA;b:CGAAGCATAACGATCGCTCCGT GCAGTAACC;c:CTGCTATCGACTGTGGAGGTACACCCTCATC;d:ATTCTCGTTCAAT GGAAGATAGCGATTTCGT;e:GGACGGCTTACCGGCTGTTCGTTTTGACGTA). (D) The 1D protein diffusion coefficients *vs* the ruggedness 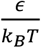. The curve is shown generically (Eq15), while the numerically obtained values of the *ruggedness* for each 30-bp DNA a-e from (C) are also shown. Notable local energy minima on the diffusion free energy landscape bring large ruggedness *ϵ*.

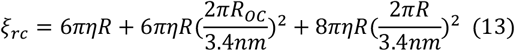

where the first term accounts for the protein (with radius *R*) translational displacement, the second term (with *R*_*OC*_ the distance between DNA and protein COM) for the curvilinear motion, and the third for the protein spinning (50). According to the Einstein’s relation

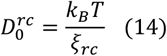

and the ultimate effective diffusion coefficient

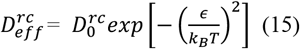

As a result, the rotation-coupled 1D protein-DNA diffusion model leads to 10 to10^2^-fold reduction in the diffusion coefficient if *R >* 1*nm* (**FIG 3B**). For example, for the spherical protein with *R* = 1.5*nm*, the diffusion coefficient for the pure translational sliding is 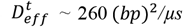 and the corrected diffusion coefficient for the rotation-coupled protein sliding is about 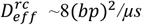, i.e., protein searches up to ∼4 *bp* per *µs*, close to the TF protein diffusion rates measured experimentally (49) (**FIG 3B**). Note that under the geometrical effects due to the protein size and DNA helical path corrections etc., the diffusion coefficients are effectively reduced by 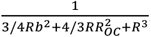, where 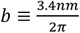 is the helical pitch of B-form DNA. Additionally, with the protein-DNA electrostatic and HB interactions modeled here, the diffusion coefficient can be further reduced, i.e., via the ruggedness *ϵ* (Eq 12), though such as a reduction is often less than 10-fold.

From generical impacts of the ruggedness 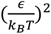. on the protein 1D diffusion rates or the diffusion free energy landscape along DNA (**FIG 3C**), one can see that *ϵ*∼1.5 *k*_*b*_*T* slightly above thermal fluctuations lead to a reduction of the local diffusion coefficient ∼ 5 times. Hence, the free energy *ruggedness* marginally above the thermal fluctuation seems to impose detectable impacts on the protein 1D diffusion rates. If one defines the ruggedness as a local feature, i.e., as being obtained numerically for the 30-bp random DNAs (**FIG 3C&D**), one can see that 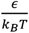 quantities fit well with Eq 15. The results confirm first that the highly non-specific protein-DNA interactions (as for DNA sequence a&b) attribute to a thermal fluctuation level of free energy ruggedness (∼1 *k*_*B*_*T*); intermediately low free energy minima (c&d to e), corresponding to consensus DNA binding sites, bring about larger values of *ϵ* (∼1.5 *k*_*B*_*T*) or lead to more rugged free energy landscape, effectively by reducing the 1D diffusion rates of the protein in search of the local DNA segment.

#### 2. Sequence-dependent collective HB dynamics at the protein-DNA interface dictate the ruggedness of the protein diffusion landscape and stepping at > 1-bp steps

To locate the protein positioning according to the interfacial HBs along DNA, we employed a measure of the collective HBs by averaging individual HB contacts at the protein-DNA interface, 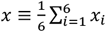,where *x*_*i*_ stands for the position of the *i*-th HB interaction site on the DNA. The free energy profile associated with *x* accordingly determines how hard the collective HB contacts shift (i.e., broken, synchronize, and reform) as the protein slides along DNA. In **FIG 4A**, the diffusion free energy profile according to the collective HB dynamics for the protein stepping along homogeneous Poly-A sequences is shown, which demonstrates 1-bp periodicity and barriers at the thermal fluctuation level, i.e., with the barrier height Δ*h*∼1*k*_*B*_*T*. A representative path consisting of 6 configurations of the protein-DNA interfacial HBs is also shown for one cycle, i.e., with 1-bp displacements for all HB contacts toward the end of the cycle. The most probable conformation ‘1’ is the one with the minimal free energy (**FIG 4A** *top*). The 1-bp stepping time statistics along poly-A DNA is obtained from the Langevin dynamics simulations and shows an exponential distribution for the stepping time regime *t <* 4 *×* 10^3^ step sampled (**FIG 4A** *bottom*).

**FIG. 4:**
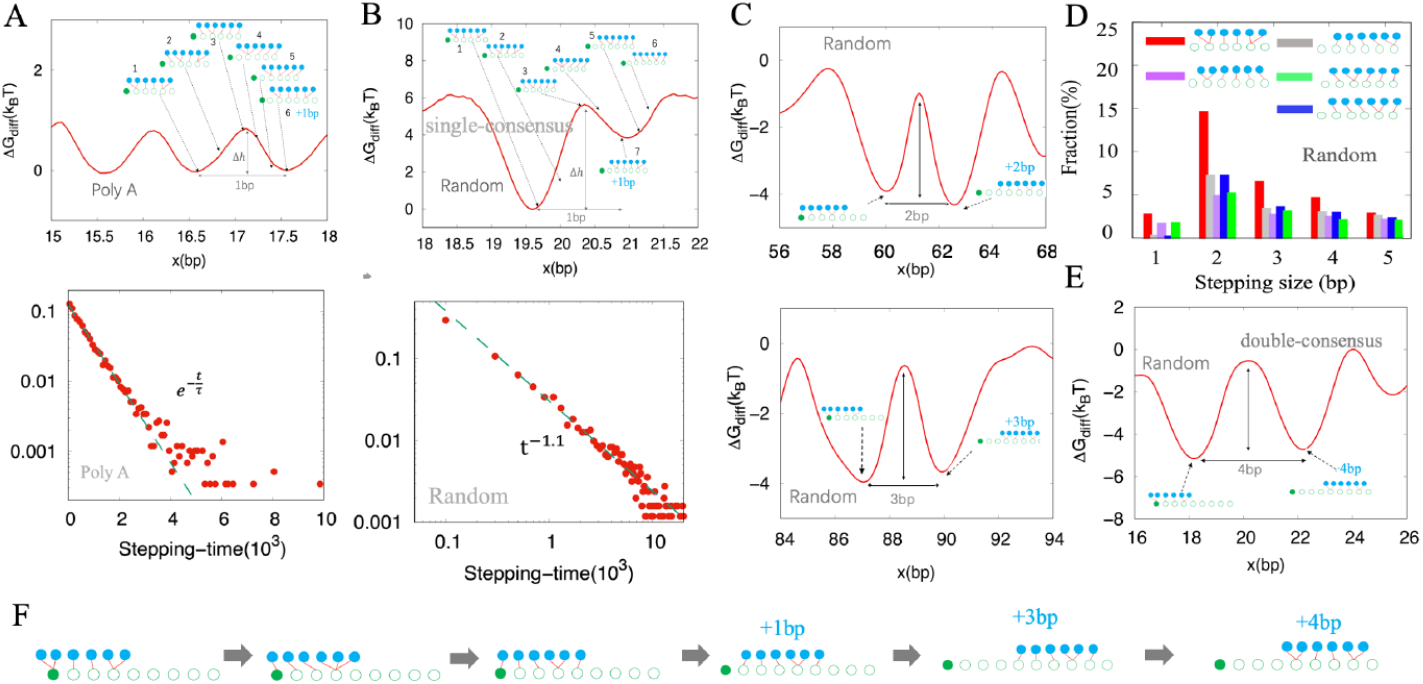
Collective hydrogen-bond (HB) dynamics at the protein-DNA interface and protein stepping statistics on DNA. (A) Diffusion free energetics according to the collective HB dynamics for protein sliding along homogeneous poly-A DNA (*top*). Multiple protein-DNA HB configurations are shown sequentially along a typical path of protein stepping in 1-bp cycle. The energy barriers are measured with height Δ*h*. The duration time for the 1-bp cycles of collective HBs as protein stepping along poly-A DNA (*bottom*). The distribution follows an exponential decay (*τ* = 3.6 × 10^3^ steps) for the short time regime (*t <* 4 *×* 10^3^ steps). (B) Diffusion free energetics according to the collective HB dynamics for protein sliding along a segment of random sequence DNA containing one single consensus site (the free energy minimum; *top*). The duration time for protein stepping 1-bp quasi-cycle on a long random DNA of 3000 bp (*bottom*). The 1-bp stepping time for the quasi-cycle obeys the power law (dashed line: *t* ^−1.1^). (C) Examples of 2-bp (*top*) and 3-bp (*bottom*) stepping sizes the random DNA sequence. The stepping sizes are consistent with the separations of the neighboring free energy minima. (D) Histogram for the fractions of the populations of the protein stepping sizes, i.e., sampled according to the collective HB configurations at the protein-DNA interface. The population for stepping along the poly-A was sampled from 0.2 *ms* trajectories, and that for the random sequence was sampled from 2 *ms* trajectory in total (0.1ms trajectory along random sequences of 150 bp each, and 20 different sequences were used). (E) An example of a short segment of random DNA sequence containing double consensus sites, offering 4-bp stepping. (F) A typical path of the HB configurations for the 4-bp stepping between the free energy minimum at *x* = 18 to that at *x* = 22 in (E).

For random DNA sequences containing single or double consensus binding cites of the protein, locally, though the DNA sequences do not repeat, 1-bp or larger than 1-bp stepping per quasi-cycle can still be identified, i.e., according to distance between neighboring free energy minima (**FIG 4B&C**). For 1-bp quasi-cycles or steps on the random DNA, one representative case is shown with 7 collective HB configurations (**FIG 4B** *top*). The stepping time statistics for various 1-bp quasi-cycles on the random sequence demonstrate a power law distribution (**Fig 4B** *bottom*), which reflects miscellaneous time scales due to various stepping barriers along the random DNA sequences (see **S3&4**).

Notably, one can identify 2-3 bp steps per quasi-cycle along the random DNA sequences (**FIG 4C**), according to distances between neighboring local energy minima (or consensus sequences of fairly low energetics), along the diffusion free energy profile of the collective HBs. The distribution of various stepping size (1-5 bp) sampled along the random DNA sequences are shown (**FIG 4D**), with a representative case for 4-bp stepping additionally demonstrated (**FIG 4E-F**). Evidently, the collective HBs move across positions of 1-bp and 3-bp configurations but of high free energetics, until they reach at the 4-bp distance collectively as a next free energy minimum (**FIG 4F**). The corresponding free energy barriers Δ*h* for the demonstrated cases of 2-bp, 3-bp and 4-bp stepping of collective HBs along the random DNA sequences range from ∼3 k_*b*_T to 4 kBT and 5 kBT (**FIG 4C & 4D** *bottom*). It is interesting to notice that the 1-bp stepping sampled along random sequences contributes only a small portion (**FIG 4D** *top*), which shows a stepping barrier ∼6 k_*b*_T in one representative case (**FIG 4B**), in contrast with the 1-bp stepping with thermal barriers (∼1 k_*b*_T) along poly-A DNA (**FIG 4A**).

Additionally, following Eq 15, the 1D effective diffusion constants are measured at ∼ 90*bp*^2^/*μs* and 30*bp*^2^/*μs* for the protein diffusing along Poly A and the above random sequences, respectively. The tunable charge of the protein is set at *Q*_*P,I*_ =+0.55e. For comparison, *Q*_*P,I*_=+0.7e is used in **FIG 3**, which results in smaller diffusion constants.

#### 3. Protein dissociation from a high-affinity DNA binding site is proceeded by diffusion to the flanking DNA sites, which always influence on the binding affinity measurements

The protein-DNA molecular interaction potential (2D PMF) *U*(*x, z*) dictates the protein diffusion and association/dissociation along DNA. Accordingly, sampling the Langevin stochastic dynamics over tens to hundreds of microseconds numerically for the protein starting at a certain DNA binding site lead to population distributions of the protein *n*(*x, z*) for diffusion or dissociation on the *x* − *z* plane, which further bring about the potential *U*(*x, z*) = −*ln*[*n*(*x, z*)*/n*_0_] (**FIG 5A-C**), where *n*_0_ is a reference population value. Consequently, the basins on the potential *U*(*x, z*) are the stabilized interaction sites with specific (e.g. TTGACT currently) or consensus core sequences at the center (at the *x*=7^th^ *bp)*. From the protein dynamics on the 2D free energy surface, one can identify distinctive representative paths for spontaneous dissociations from DNA (i.e., out of the electrostatic zone), e.g., starting from the specific DNA binding site (**FIG 5A** *top*). The dominant paths for the protein dissociations are those first via diffusion to the *flanking* non-specific DNA binding sites, and then with protein dissociations from the flanking DNA (∼93.5%). The straightforward paths for the protein dissociations ‘vertically’ (along *z*) to the DNA length account for only a small percentile (∼3%) due to high energy barriers (see **FIG S10**). Additionally, there are ‘diffusion-coupled’ paths (i.e., ‘tilted’ in between x and z directions towards the outer space; ∼3.5%; **FIG 5A** *top* and **FIG 5D**), The histogram (**FIG5A** *bottom*) shows the dwell time distribution as the protein remains bound to DNA, i.e., within the electrostatic zone. The mean protein dissociation time is ∼ 0.25 ms, starting from the specific DNA binding site as the 6-bp center core on the 15-bp DNA.

**FIG. 5:**
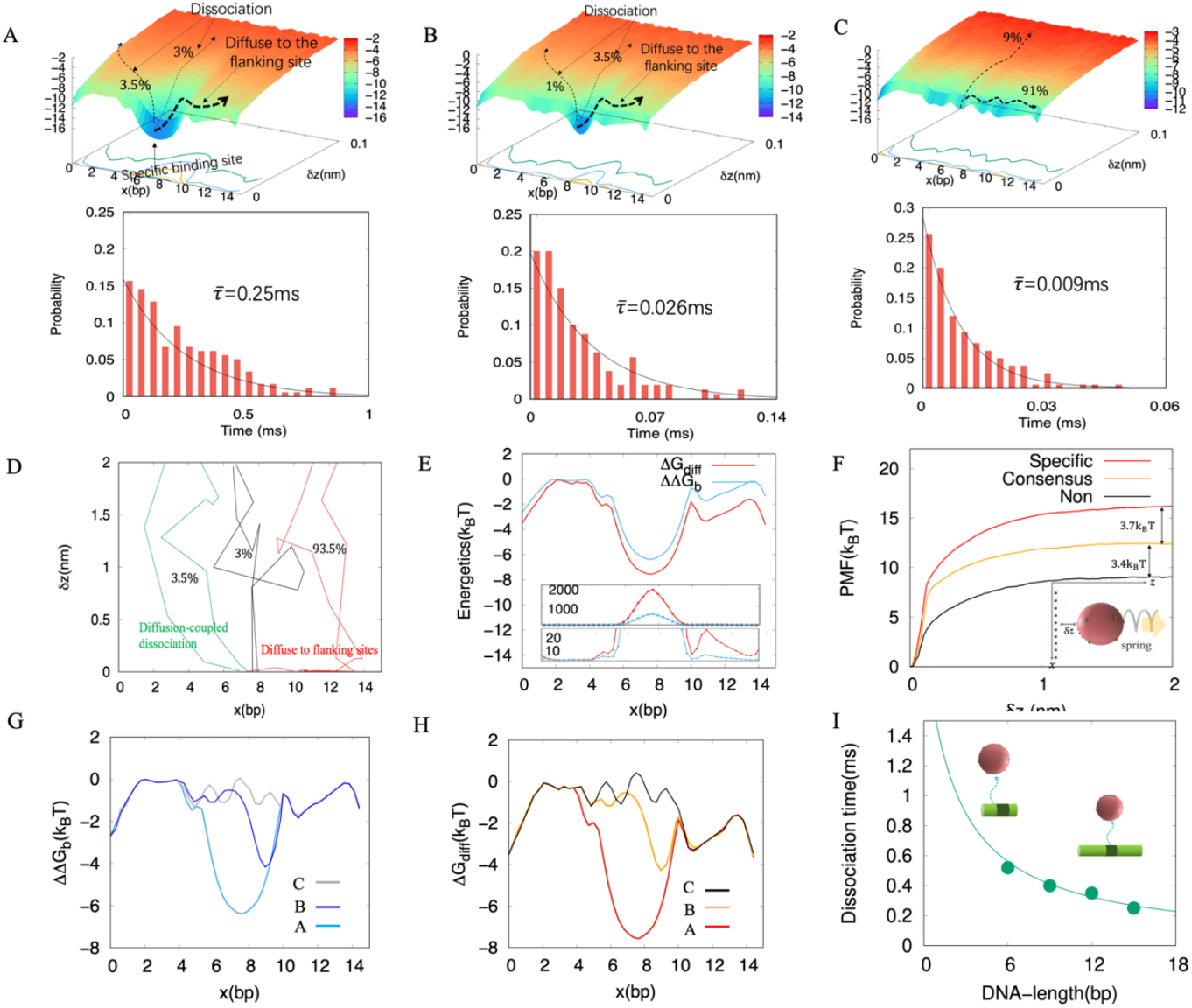
Protein dissociation vs diffusion along DNA from the Langevin dynamics simulations. (A) The 2D PMF *U*(*x, z*) ∼ −*ln*[*n*(*x, z*)] obtained numerically from the populations *n*(*x, z*) of the protein, sampled at the protein-DNA interface (*top*), with the protein initially bound to the specific DNA (TTGACT). Protein dominantly diffuses to the flanking nonspecific site and then dissociates, occasionally also dissociates in a diffusion-coupled manner (the *titled* path), or dissociate directly (*vertically* along z) from the DNA. Histogram plotted for the bound population (*bottom*), showing the dwell time distribution of the protein remaining bound to DNA. The mean dwell or dissociation time is ∼ 0.25 ms. (B) *U*(*x, z*) obtained from the population of protein sampled around a consensus sequence (TTCACT). Histogram on the dwell time (*bottom*) and the mean dissociation time ∼0.026 ms. (C) *U*(*x, z*) obtained from the population of protein sampled starting from a nonspecific DNA core sequence (TTCTCT). Histogram on the dwell time (*bottom*) and the mean dissociation time ∼0.01ms. (D) Three typical paths sampled from simulations from (A) representing the diffusion-coupled (*titled* path, green), direct (*vertical* path, black) dissociation, and diffusion-dominated path (*red*). The corresponding population percentages were calculated from 200 sampled dissociation events. (E) 1D diffusion free energy landscape Δ *G*_*diff*_ (*x*) obtained from the Langevin dynamics simulations for (A) according to the protein populations along DNA, with protein-DNA HB contacts maintained for the diffusion; shown as well the relative association free energy profile ΔΔ*G*_*b*_(*x*) along DNA, for populations of protein in association with DNA within the electrostatic zone. The minimum of Δ*G*_*diff*_(*x*) is ∼ 1.2 *k*_*B*_T deeper than that of ΔΔ *G*_*b*_ (*x*) in this case. The inset shows the corresponding protein populations for diffusion vs association or binding on DNA. (F) The 1D PMFs by pulling protein quasi-statically (along z) away from DNA, from the specific, consensus and non-specific sites, where the protein is bound initially (as from A, B, and C). (G) and (H) show the free energetics ΔΔ*G*_*b*_(*x*) and Δ*G*_*diff*_ (x), respectively, for the protein along the short DNA sequences (15 bp length; as from A, B and C). (I) The predicated DNA length dependency of the measured protein dissociation time (1/rate), with the protein initially bound at the specific (from A) or a high-affinity DNA binding site. The fitting curves are 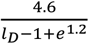 (see **S5**).

Similarly, the 2D potential *U*(*x, z*) centered around a consensus DNA binding site (TTGACT) is shown in **FIG 5B** (*top*). The dissociations have slightly more *vertical* paths (∼3.5%) than the *tilted* diffusion-coupled paths (∼1%), likely due to lower vertical dissociation barrier of the consensus site than the specific site. Still, the dominant paths are those via the diffusion to the flanking site first (∼95.5%). Overall, the mean protein dissociation time from the 15-bp DNA is about 0.026 *ms*, much faster than that from the DNA with the specific core. In further comparison, the 2D potential *U*(*x, z*) centered around a non-specific core sequence (TTCTCT) is shown in **FIG 5C** (*top*). In this case, ∼ 9% of the population of the protein dissociation happens at the core site, or say, ∼91% population continues for the diffusion. The mean dissociation time is ∼0.01 *ms*. Note that only for the pure non-specific DNA (with the core non-specific binding site), the dwell time obeys well an exponential distribution (**FIG 5C** *bottom*), which indicates single-rate events. In contrast, the protein dissociation from the DNA containing the specific or consensus core site at the center includes multiple kinetic events of mixed rates, hence, the dwell-time distributions deviate notably from the exponential form (**FIG 5A&B** *bottom*). If one however still uses the average dwell time to estimate the dissociation rate 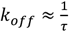, and further estimates binding the free energy difference between two core sites (∼6 bp) based on the dissociation events from the full-length DNA construct A & B (15 bp each) as 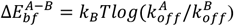, assuming binding rates are identical or independent of DNA sequences, then we estimate 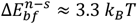 between the non-specific and specific DNA, 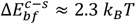 between the consensus and specific DNA, and 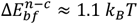 between the nonspecific and consensus DNA. Comparing with the free energy minima differences revealed from1D ΔΔ*G*_*b*_(*x*) and Δ*G*_*diff*_(*x*) (**Table 1**), however, one sees notable under-estimation of the free energy differences via the above approximations, in particular, relative to the non-specific DNA. Similar issues were identified recently, via the atomic MD simulations pulling protein away from DNA to measure the relative binding free energetics between the specific and non-specific DNA binding sites (51).

**Table 1:** Comparing relative protein-DNA binding free energetics obtained from different approaches.

| The *relative* binding free energetic differences quantified via various approaches | $\Delta E_b^{n-s}$ (non-specific vs specific) | $\Delta E_b^{c-s}$ (consensus vs specific) | $\Delta E_b^{n-c}$ (non-specific vs consensus) |
| --- | --- | --- | --- |
| (i) Via estimations on $k_{off}$ (with flanking sequences)<br>$\Delta E_{bf}$ | 3.3 kBT | 2.3 kBT | 1.1 kBT |
| (ii) Via the relative binding free energy profile<br>$\Delta\Delta G_b(x)$: $\Delta E_{bb}$ | 5.8 kBT | 2.3 kBT | 3.5 kBT |
| (iii) Via the diffusion free energy landscape<br>$\Delta G_{diff}(x)$: $\Delta E_{bd}$ (or pulling protein from DNA) | 7.1 kBT | 3.3 kBT | 3.8 kBT |

One can further map the numerically constructed *U*(*x, z*) to 1D along DNA (**FIG 5E**), for the protein diffusion and association profiling as demonstrated analytically (**FIG 2C** *bottom*) for Δ *G*_*diff*_(*x*) and ΔΔ *G*_*b*_(*x*), respectively. One confirms that via the stochastic Langevin dynamics, as the protein rotation DOFs are not constrained but numerically integrated with the translational DOFs, the relative protein association profiling ΔΔ*G*_*b*_(*x*) smoothens over the diffusion landscape Δ*G*_*diff*_(*x*)○ As the 2D potential surface **FIG 5A** maps to **FIG 5E**, the minimum of Δ*G*_*diff*_(*x*) shows ∼ 1.2*k*_*B*_T deeper than that of ΔΔ*G*_*b*_ (*x*), and the corresponding population peak on the diffusion landscape is about twice steeper than that on the association profiling. The trend maintains for the 1D profiling mapped from *U*(*x, z*) around the consensus and non-specific core sites (from **FIG 5B&C** to **FIG 5G&H**), though less prominent as that in the specific binding case. One can also compare the *relative* binding free energetics between the core DNA binding sequences (specific vs consensus vs non-specific), i.e., revealed at the free energy *minima* of ΔΔ*G*_*b*_ (*x*) (**FIG 5G**) and Δ*G*_*diff*_(*x*) (**FIG 5H**). The results show that the readouts from ΔΔ*G*_*J*_ (*x*) are always smaller than that from Δ*G*_*diff*_(*x*), with 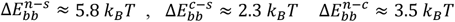 from ΔΔ *G*_*b*_ (*x*) minima and 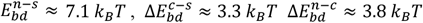 from Δ*G*_*diff*_(*x*) minima (see **Table 1**). It is because Δ*G*_*diff*_(*x*) reflects precisely the sequence-dependent energetics (via the HB contact) at each bp position, while ΔΔ*G*_*b*_ (*x*) integrates additional DOFs (e.g., the protein rotation) as the protein bound comparatively loosely within the electrostatic zone. The precise energetics differences between free energy minima on Δ*G*_*diff*_(*x*) can also be revealed by pulling the protein initially bound at the core site of DNA (specific vs consensus vs non-specific) away from DNA, quasi-statically, and measuring the work or free energetics (**FIG 5F**).

As protein diffusion along DNA are intervened by protein dissociation events from DNA, which happen relatively infrequently, the length of the DNA segment to which the protein is bound affects the dissociation time or rates measured on it. Via the Langevin dynamics simulations, **FIG 5I** shows, interestingly, that the longer the DNA length *l*_*D*_, the shorter the mean dissociation time or dwell time of protein in association with the DNA. Essentially, for a DNA construct with a high-affinity DNA binding site located at the center and non-specific sites flanking the surround region, the longer DNA supports higher chances of the protein to diffuse to the flanking sites to dissociate, and the shorter DNA allows more chances of the protein to diffuse back to the center site to be trapped. The quantitative relation between the mean dissociation time and *l*_*D*_ can be further derived 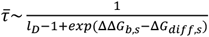(see **FIG S11**), as shown in the **FIG 5I**. Again, one can see that the subtle difference between ΔΔ*G*_*b*_ (*x*) and Δ*G*_*diff*_(*x*) around the specific binding site can be exponentially amplified to population differences and detectable dissociation time for measurements.

## IV. DISCUSSION

In current work,we have constructed a minimal structure-based physical model of protein diffusion and association-dissociation along DNA to illustrate the protein target search dynamics and the corresponding free energy landscape, in particular, around the target or consensus binding site surrounded by flanking DNA sequences. As protein diffuses along 1D DNA about tens of bps distances at sub-millisecond time intervals, and occasionally dissociates from DNA into 3D nucleus space around millisecond time scale, there exists a timescale separation between the protein 1D diffusion and dissociation. Upon construction of the 2D free energy surface of the protein diffusion-association around the binding site, we further map the 2D surface to 1D along DNA to account for diffusion and association-dissociation dynamics at disparate time scales. We envision that the protein 1D diffusional free energy landscape can be constructed within milliseconds but over microseconds, as protein intensively scans DNA sequences to search and locate the specific site. Such processes can be detected presumably from single molecule measurements, though it remains challenging for detecting individual molecular motoions at both high temporal (submillisecond) and spatial (subnanometer) resolutions, simultaneously (11, 52). Meanwhile, the protein binding or functional association profiling can be measured on DNA segments via highthroughput experimental methods such as the PBM (protein binding microarray) (53-55), or genomewide *in vivo* via ChIP-seq or high-resolutoin ChIP-exo technologies (56-58). Consequently, one anticipates that the relative protein association profiling mapped along 1D DNA in our simplied phyiscal representation can be made comparable with the high-resolution datasets genomewide.

In current model, a spherical protein is adopted with polarized surface charges, effectively, interacting with linear DNA containing negative charges and with helical geometry. The model explicitly accounts for protein-DNA electrostatic interactions, generically, along with the implicit solvent and ionic screening effects. Importantly, we also modeled HB interactions or contacts at the protein-DNA interface, which are orientation-dependent and sensitive to DNA sequences. The orientation is particularly impacted by the protein rotation in current model, and is fully taken into consideration along with the protein translation. The DNA sequence dependency is presumably encoded by the strengths of the interfacial HBs. Technically, we incorporate the sequence dependency into the interfacial HBs via the PWM (position weight matrix), commonly used in bioinformatics to well account for the favored DNA binding motifs by individual TF proteins (59, 60). With the protein-DNA electrostatic and HB interactions explicitly modeled, along with the DNA helical path corrections for hydrodynamics, our model well captures the time scale of the protein diffusion at microseconds or sub-milliseconds, via the Langevin stochastic dynamics simulation. The diffusional coefficients obtained numerically are consistent at the order of magnitude with experimental measurements (49, 61, 62), without additional parameter fitting or tuning other than setting up the effective charges and size of the protein. Upon the full characterization of the protein 1D diffusion free energy landscape, one captures non-trivial ruggedness of the free energy landscape above thermal fluctuation level, in particular, for DNA with notable local free energy minima or barriers, which serve for consensus of the target DNA sequences.

Once the protein diffusion rates or timescales along DNA are well calibrated, we were able to use current model to investigate the sequence-dependent protein sliding/stepping dynamics along DNA, with much higher computational efficiency for sufficient statistics than performing structure-based simulations, in particular, all-atom molecular dynamics (MD) simulations, which are computationally demanding (63, 64). Essentially, we demonstrate that the sequence-dependent collective HB dynamics at the protein-DNA interface well dictates the protein diffusion barriers or ruggedness, as well as the protein stepping dwell times and patterns. As seen from recent atomic MD studies (17, 51), the interfacial HBs are constantly formed between the charged (Arg and Lys) or polar protein residues and the DNA backbone atoms, or occasionally with DNA bases at the specific site. In current minimal structure-based model, we additionally noticed that the protein stepping sizes (i.e., the number of bps for the interfacial collective HBs to reset or between two neighboring diffusion free energy minima) vary stochastically along DNA, and the distribution of the stepping sizes (e.g. >1bp) distinguish between different DNA sequence patterns, e.g., homogeneous poly-A (dominantly 1-bp) vs various random DNA sequences (more distributions to >1bp). The uniform energy barriers for the collective HB contacts along the homogeneous DNA sequences lead to an exponential distribution of the stepping time, while the diverse energy barriers for stepping along the random sequence give a power law time distribution, indicating multiple transition timescales involved. Additionally, synchronizing the collective HB dynamics appears much easier (low barriers or taking shorter time) for protein moving along the homogeneous sequences than along the random DNA sequences of variable features. Accordingly, the diffusion rate of protein along the homogeneous sequences is to be larger than that on the random sequences with various features or increased ruggedness. Such predictions on the sequence dependent stepping patterns of the protein can be tested experimentally, e.g., by designing periodic patterns of the DNA or utilizing short repetitive sequences (65, 66), and measuring the corresponding protein diffusion rates and patterns, correspondingly.

As an extension of protein diffusional search dynamics to millisecond time scale, spontaneous protein dissociations from DNA can be well sampled in the Langevin dynamics simulation. We in particular define the protein escaping from the protein-DNA electrostatic zone (*δ*z > 2 - 3nm) as the dissociation (or micro-dissociation). The spontaneous protein dissociation paths simulated starting from a high-affinity DNA binding site fall into three categories or populations, with the dominant population (> 95%) diffusing to flanking DNA sites first and then dissociating from the non-specific flanking sites. Rarely there are protein populations dissociating directly or ‘vertically’ from the initial binding site, due to the steep free energy barriers following such paths from the high-affinity DNA binding site. Consequently, impacts from the non-specific flanking sites would become highly pronounced for measuring binding profiling or relative binding affinities of the protein to the full-length DNA containing flanking sites surrounding the initial core binding site. That says, the DNA construct length impacts significantly the apparent measured dissociate rates of the protein or further the relative binding affinities. Furthermore, one may intuitively consider the longer DNA leading to the longer dissociation time, or lower dissociation rates. However, our simulations show that the longer DNA containing a high-affinity DNA core binding site results in the shorter mean dissociation time or higher dissociation rates. The longer DNA essentially supports the protein to diffuse extensively to the non-specific flanking sites to increase the chances of dissociation, rather than allows the protein to move back to the initial high-affinity binding site to be trapped again. Hence, by exploring the protein dissociation from the DNA construct, we show notably the DNA length effects and the flanking sequence impacts to the measured dissociation rates, based on our simple physical model of protein-DNA interactions and stochastic dynamics. The working scenarios here on the flanking sequence impacts offer physical insight to the data-driven bioinformatic or genome-wide mapping studies (67-69). Note that currently the protein dissociation is defined above DNA over 2-3 nm distance, depending on the solution salt concentrations. A so-called macro-dissociation at least beyond ∼30 nm (or 100 bp) was considered previously (8, 70), e.g., as the protein having larger chance to be away than returning to DNA. The macro-dissociation involves larger length scale beyond the persistence length of DNA (∼150 bp), hence DNA coiling and related dynamics become important, which is beyond our current model of fairly short local DNA (tens of bps) in a static and linear form.

Notably, via current model, we constructed generically the 2D molecular potential or the free energy surface of the protein diffusion-association. By further mapping the 2D potential surface to 1D, we have demonstrated both analytically and numerically that the 1D relative protein association free energy profiling ΔΔ*G*_*b*_ (*x*) along DNA, i.e., due to populations of protein bound to DNA within the electrostatic zone (*δ*z < 2 - 3nm), smoothens over the 1D protein diffusion free energy landscape Δ*G*_*diff*_ (*x*), as the protein slides along DNA for sequence detection by maintaining interfacial HB contacts (*δ*z<0.2 nm). Such short-range HB interactions at the protein-DNA interface reflect precisely the protein readout on the DNA sequences and also restrain the protein conformational DOFs such as rotations modeled in current study. In comparison, the long-range protein-DNA electrostatics interactions remain fairly uniform along DNA in current model, which ensure protein-DNA association with dynamical DOFs of protein softly constrained or accommodated. The protein bound to DNA electrostatically can thus undergo partial rotation and spin, along with conformational changes and micro-hopping (not yet addressed in current model), as the protein-DNA interfacial HB contacts frequently or largely break. Among these flexibilities, we particularly considered the protein spatial rotation and spin in current model as the extra DOFs. Numerically our simulations show that partial protein rotations (up to 20-40 degrees) are allowed or relaxed, when protein remains bound within the electrostatic zone of DNA but breaks majorities of the protein-DNA HB contacts. Consequently, the protein binding profiling along DNA accommodating the extra DOFs smoothens over the protein 1D diffusion free energy landscape, which is highly rugged to reflect closely the DNA sequence variations. Though the smoothen effect reveals only 1-2 k_*b*_T in current model, it already causes multiple times of population differences (exp(2.3)∼10), predicted between the protein binding and diffusion free energy minima or population peaks along DNA. In reality, one expects that the smoothen effects become more pronounced when protein conformational fluctuations are further accounted for. Future single-molecule experimental measurements with sufficient spatial and temporal resolutions may be able to resolve the stepwise dynamics of the protein along DNA, and characterize the protein diffusion free energy landscape directly, allowing comparison with high-resolution protein association profiling measured from *in-vitro* assay, so that to validate our model predictions.

Furthermore, protein can re-associate with DNA after dissociation. It was recently suggested that sequence specificity of DNA binding is dominated by protein association (71). Nevertheless, protein re-association after initial association and dissociation need to be further considered. The re-association events were occasionally captured in our simulations, but the dynamics have not been yet examined in current work. Presumably, when protein enters or re-enters the electrostatic zone to associate with DNA, it needs to adjust its orientation to impose the positively charged surface area favorably towards the DNA. Initial binding events and re-binding events with memory effects of dissociation may adopt different adjustments during this stage. DNA sequence impacts from the flanking DNA sites, either due to DNA shape effects (72, 73) or 1D diffusion suggested here and a recent work (51), can play additional roles in the association and re-association. Furthermore, as the protein binds tightly to the DNA specific or consensus site within the HB interaction range, refined adjustments of the protein orientation DOFs are expected to take place to support formations of collective HBs at the protein-DNA interface. Such adjustments are likely highly depend on DNA sequences locally. Still, how the DNA sequences impact on the close associations and re-associations of protein and DNA remain to be further investigated.

## Acknowledgements

JY has been supported by UC-CRCC (Cancer Research Coordinating Committee) grant C23CR5636. BW is supported by NSFC (National Natural Science Foundation of China) grant #12304246.

## References

1. F. Cozzolino, I. Iacobucci, V. Monaco, M. Monti, Protein-DNA/RNA Interactions: An Overview of Investigation Methods in the-Omics Era. J Proteome Res 20, 3018–3030 (2021).

2. D. M. Suter, Transcription Factors and DNA Play Hide and Seek. Trends Cell Biol 30, 491–500 (2020).

3. A. A. Shvets, M. P. Kochugaeva, A. B. Kolomeisky, Mechanisms of Protein Search for Targets on DNA: Theoretical Insights. Molecules 23 (2018).

4. J. M. Franco-Zorrilla et al., DNA-binding specificities of plant transcription factors and their potential to define target genes. Proc Natl Acad Sci U S A 111, 2367–2372 (2014).

5. A. Tafvizi, F. Huang, A. R. Fersht, L. A. Mirny, A. M. van Oijen, A single-molecule characterization of p53 search on DNA. Proc Natl Acad Sci U S A 108, 563–568 (2011).

6. P. H. von Hippel, O. G. Berg, Facilitated target location in biological systems. Journal of Biological Chemistry 264, 675–678 (1989).

7. R. F. Bruinsma (2002) Physics of Protein-DNA Interaction. in Physics of bio-molecules and cells. Physique des biomolécules et des cellules, eds F. Flyvbjerg, F. Jülicher, P. Ormos, F. David (Springer Berlin Heidelberg, Berlin, Heidelberg), pp 1–68.

8. S. E. Halford, J. F. Marko, How do site-specific DNA-binding proteins find their targets? Nucleic Acids Research 32, 3040–3052 (2004).

9. P. Hammer. et al., The lac Repressor Displays Facilitated Diffusion in Living Cells Science 336 (2012).

10. P. Hammar et al., The lac repressor displays facilitated diffusion in living cells. Science 336, 1595–1598 (2012).

11. J. Elf, I. Barkefors, Single-Molecule Kinetics in Living Cells. Annual Review of Biochemistry 88, 635–659 (2019).

12. I. Izeddin et al., Single-molecule tracking in live cells reveals distinct target-search strategies of transcription factors in the nucleus. eLife 3, e02230 (2014).

13. C. Monico, M. Capitanio, G. Belcastro, F. Vanzi, F. S. Pavone, Optical Methods to Study Protein-DNA Interactions in Vitro and in Living Cells at the Single-Molecule Level. International Journal of Molecular Sciences 14 (2013).

14. X. S. Xie, P. J. Choi, G.-W. Li, N. K. Lee, G. Lia, Single-Molecule Approach to Molecular Biology in Living Bacterial Cells. Annual Review of Biophysics 37, 417–444 (2008).

15. Y. M. Wang, R. H. Austin, E. C. Cox, Single molecule measurements of repressor protein 1D diffusion on DNA. Phys Rev Lett 97, 048302 (2006).

16. J. Yoo, D. Winogradoff, A. Aksimentiev, Molecular dynamics simulations of DNA-DNA and DNA-protein interactions. Curr Opin Struct Biol 64, 88–96 (2020).

17. L. Dai, Y. Xu, Z. Du, X. D. Su, J. Yu, Revealing atomic-scale molecular diffusion of a plant-transcription factor WRKY domain protein along DNA. Proc Natl Acad Sci U S A 118 (2021).

18. M. R. Tucker, S. Piana, D. Tan, M. V. LeVine, D. E. Shaw, Development of Force Field Parameters for the Simulation of Single- and Double-Stranded DNA Molecules and DNA-Protein Complexes. J Phys Chem B 126, 4442–4457 (2022).

19. C. Tan, S. Takada, Dynamic and Structural Modeling of the Specificity in Protein-DNA Interactions Guided by Binding Assay and Structure Data. J Chem Theory Comput 14, 3877–3889 (2018).

20. L. Dai, J. Yu, Inchworm stepping of Myc-Max heterodimer protein diffusion along DNA. Biochemical and Biophysical Research Communications 533, 97–103 (2020).

21. L. S. Bigman, Y. Levy, Protein Diffusion on Charged Biopolymers: DNA versus Microtubule. Biophys J 118, 3008–3018 (2020).

22. P. Dey, A. Bhattacherjee, Structural Basis of Enhanced Facilitated Diffusion of DNA-Binding Protein in Crowded Cellular Milieu. Biophysical Journal 118, 505–517 (2020).

23. U. Gerland, J. D. Moroz, T. Hwa, Physical constraints and functional characteristics of transcription factor-DNA interaction. Proc Natl Acad Sci U S A 99, 12015–12020 (2002).

24. A. R. Dinner, A. Sali, L. J. Smith, C. M. Dobson, M. Karplus, Understanding protein folding via free-energy surfaces from theory and experiment. Trends Biochem Sci 25, 331–339 (2000).

25. D. J. Wales, T. V. Bogdan, Potential energy and free energy landscapes. J Phys Chem B 110, 20765–20776 (2006).

26. P. Das, M. Moll, H. Stamati, L. E. Kavraki, C. Clementi, Low-dimensional, free-energy landscapes of protein-folding reactions by nonlinear dimensionality reduction. Proc Natl Acad Sci U S A 103, 9885–9890 (2006).

27. P. G. Wolynes, Evolution, energy landscapes and the paradoxes of protein folding. Biochimie 119, 218–230 (2015).

28. F. Pietrucci, Strategies for the exploration of free energy landscapes: Unity in diversity and challenges ahead. Reviews in Physics 2, 32–45 (2017).

29. D. Wang et al., Efficient sampling of high-dimensional free energy landscapes using adaptive reinforced dynamics. Nat Comput Sci 2, 20–29 (2022).

30. M. Slutsky, L. A. Mirny, Kinetics of Protein-DNA Interaction: Facilitated Target Location in Sequence-Dependent Potential. Biophysical Journal 87, 4021–4035 (2004).

31. L. Mirny et al., How a protein searches for its site on DNA: the mechanism of facilitated diffusion. Journal of Physics A: Mathematical and Theoretical 42, 434013 (2009).

32. I. Leven, Y. Levy, Quantifying the two-state facilitated diffusion model of protein–DNA interactions. Nucleic Acids Research 47, 5530–5538 (2019).

33. P. C. Blainey, A. M. van Oijen, A. Banerjee, G. L. Verdine, X. S. Xie, A base-excision DNA-repair protein finds intrahelical lesion bases by fast sliding in contact with DNA. Proc Natl Acad Sci U S A 103, 5752–5757 (2006).

34. M. F. Berger, M. L. Bulyk, Universal protein-binding microarrays for the comprehensive characterization of the DNA-binding specificities of transcription factors. Nat Protoc 4, 393–411 (2009).

35. S. W. Ho, G. Jona, C. T. Chen, M. Johnston, M. Snyder, Linking DNA-binding proteins to their recognition sequences by using protein microarrays. Proc Natl Acad Sci U S A 103, 9940–9945 (2006).

36. T. S. Furey, ChIP–seq and beyond: new and improved methodologies to detect and characterize protein–DNA interactions. Nature Reviews Genetics 13, 840–852 (2012).

37. R. Jothi, S. Cuddapah, A. Barski, K. Cui, K. Zhao, Genome-wide identification of in vivo protein-DNA binding sites from ChIP-Seq data. Nucleic acids research 36, 5221–5231 (2008).

38. V. R. Yella et al., Flexibility and structure of flanking DNA impact transcription factor affinity for its core motif. Nucleic Acids Res 46, 11883–11897 (2018).

39. A. Afek, J. L. Schipper, J. Horton, R. Gordân, D. B. Lukatsky, Protein-DNA binding in the absence of specific base-pair recognition. Proc Natl Acad Sci U S A 111, 17140–17145 (2014).

40. P. Zhang et al., Deep flanking sequence engineering for efficient promoter design using DeepSEED. Nat Commun 14, 6309 (2023).

41. V. Dahirel, F. Paillusson, M. Jardat, M. Barbi, J. M. Victor, Nonspecific DNA-protein interaction: why proteins can diffuse along DNA. Phys Rev Lett 102, 228101 (2009).

42. M. Doi, Soft Matter Physics (Oxford University Press, 2013).

43. S. V. Goryainov, A model of phase transitions in double-well Morse potential. Physica B: Condensed Matter 407, 4233–4237 (2012).

44. K. Brameld, Siddharth Goddard, William A., Distance Dependent Hydrogen Bond Potentials for Nucleic Acid Base Pairs from ab Initio Quantum Mechanical Calculations (LMP2/cc-pVTZ). The Journal of Physical Chemistry B 101, 4851–4859 (1997).

45. K. Nishida, M. C. Frith, K. Nakai, Pseudocounts for transcription factor binding sites. Nucleic Acids Res 37, 939–944 (2009).

46. I. Erill, M. C. O’Neill, A reexamination of information theory-based methods for DNA-binding site identification. BMC Bioinformatics 10, 57 (2009).

47. Y. P. Xu, H. Xu, B. Wang, X. D. Su, Crystal structures of N-terminal WRKY transcription factors and DNA complexes. Protein Cell 11, 208–213 (2020).

48. R. Zwanzig, tDiffusion in a rough potential. Proc Natl Acad Sci U S A 85, 2029–2030 (1988).

49. P. C. Blainey et al., Nonspecifically bound proteins spin while diffusing along DNA. Nature Structural & Molecular Biology 16, 1224–1229 (2009).

50. B. Bagchi, P. C. Blainey, X. S. Xie, Diffusion constant of a nonspecifically bound protein undergoing curvilinear motion along DNA. J Phys Chem B 112, 6282–6284 (2008).

51. C. Al Masri, B. Wan, J. Yu, Non-specific vs specific DNA binding free energetics of a transcription factor domain protein. Biophys J (2023).

52. T. Ha, C. Kaiser, S. Myong, B. Wu, J. Xiao, Next generation single-molecule techniques: Imaging, labeling, and manipulation in vitro and in cellulo. Mol Cell 82, 304–314 (2022).

53. M. Godoy et al., Improved protein-binding microarrays for the identification of DNA-binding specificities of transcription factors. Plant J 66, 700–711 (2011).

54. K. K. Andrilenas, A. Penvose, T. Siggers, Using protein-binding microarrays to study transcription factor specificity: homologs, isoforms and complexes. Brief Funct Genomics 14, 17–29 (2015).

55. M. A. Hume, L. A. Barrera, S. S. Gisselbrecht, M. L. Bulyk, UniPROBE, update 2015: new tools and content for the online database of protein-binding microarray data on protein-DNA interactions. Nucleic Acids Res 43, D117–122 (2015).

56. Ho S. Rhee, B. F. Pugh, Comprehensive Genome-wide Protein-DNA Interactions Detected at Single-Nucleotide Resolution. Cell 147, 1408–1419 (2011).

57. P. J. Skene, S. Henikoff, A simple method for generating high-resolution maps of genome-wide protein binding. Elife 4, e09225 (2015).

58. W. J. de Jonge, M. Brok, P. Lijnzaad, P. Kemmeren, F. C. Holstege, Genome-wide off-rates reveal how DNA binding dynamics shape transcription factor function. Mol Syst Biol 16, e9885 (2020).

59. A. Boytsov, S. Abramov, V. J. Makeev, I. V. Kulakovskiy, Positional weight matrices have sufficient prediction power for analysis of noncoding variants. F1000Res 11, 33 (2022).

60. X. Ma, D. Ezer, C. Navarro, B. Adryan, Reliable scaling of position weight matrices for binding strength comparisons between transcription factors. BMC Bioinformatics 16, 265 (2015).

61. J. Gorman, E. C. Greene, Visualizing one-dimensional diffusion of proteins along DNA. Nature Structural & Molecular Biology 15, 768–774 (2008).

62. J. H. Kim, R. G. Larson, Single-molecule analysis of 1D diffusion and transcription elongation of T7 RNA polymerase along individual stretched DNA molecules. Nucleic Acids Research 35, 3848–3858 (2007).

63. J. C. Phillips et al., Scalable molecular dynamics on CPU and GPU architectures with NAMD. J Chem Phys 153, 044130 (2020).

64. D. Lu, HanChen, MohanLin, LinCar, RobertoE, WeinanJia, Weile Zhang, Linfeng, 86 PFLOPS Deep Potential Molecular Dynamics simulation of 100 million atoms with ab initio accuracy. Computer Physics Communications 259, 107624 (2021).

65. X. Liao et al., Repetitive DNA sequence detection and its role in the human genome. Commun Biol 6, 954 (2023).

66. C. A. Horton et al., Short tandem repeats bind transcription factors to tune eukaryotic gene expression. Science 381, eadd1250 (2023).

67. J. L. Stringham, A. S. Brown, R. A. Drewell, J. M. Dresch, Flanking sequence context-dependent transcription factor binding in early Drosophila development. BMC Bioinformatics 14, 298 (2013).

68. R. Gordân et al., Genomic regions flanking E-box binding sites influence DNA binding specificity of bHLH transcription factors through DNA shape. Cell Rep 3, 1093–1104 (2013).

69. S. Inukai, K. H. Kock, M. L. Bulyk, Transcription factor-DNA binding: beyond binding site motifs. Curr Opin Genet Dev 43, 110–119 (2017).

70. E. G. Marklund et al., Transcription-factor binding and sliding on DNA studied using micro- and macroscopic models. Proceedings of the National Academy of Sciences 110, 19796 (2013).

71. E. Marklund et al., Sequence specificity in DNA binding is mainly governed by association. Science 375, 442–445 (2022).

72. R. Rohs et al., The role of DNA shape in protein-DNA recognition. Nature 461, 1248–1253 (2009).

73. J. Li, T. P. Chiu, R. Rohs, Predicting DNA structure using a deep learning method. Nat Commun 15, 1243 (2024).

