## Supplementary Material for "Protein target search diffusion-association/dissociation free energy landscape around DNA binding site with flanking sequences"

Biao Wan[1] and Jin Yu[2]

[1]*Wenzhou Institute, University of Chinese Academy of Sciences, Wenzhou, China*

[2]*Department of Physics and Astronomy, Department of Chemistry, NSF-Simons Center for Multiscale Cell Fate Research, University of California, Irvine, California, United States of America*

### 1. Langevin equations of the spherical model

#### A. Translational motion

The center of mass of the protein evolves under the full potential

$$\dot{\boldsymbol{R}}_{COM} = -\frac{1}{\zeta}\nabla_{R_{COM}}H + \sqrt{\frac{2k_BT}{\zeta}}\boldsymbol{w} \qquad \text{(S1)}$$

where $\zeta = 6\pi\eta R$ is the viscous drag for the bead with radius of R, $k_B$ is the Boltzmann constant and $w$ is the Gaussian noise with zero mean and unit variance.

The resultant force exerted at the center of mass causes the translational displacement. The over-damped Langevin dynamics of the COM of the protein are then

$$x(t+dt) = x(t) - \frac{1}{\zeta}\partial_x H dt + \sqrt{\frac{2k_BTdt}{\zeta}}w$$

$$y(t+dt) = y(t) - \frac{1}{\zeta}\partial_y H dt + \sqrt{\frac{2k_BTdt}{\zeta}}w \quad (S2)$$

$$z(t+dt) = z(t) - \frac{1}{\zeta}\partial_z H dt + \sqrt{\frac{2k_BTdt}{\zeta}}w$$

where the time-step is $dt = 1ps$.

#### B. Dynamics of rotational degrees

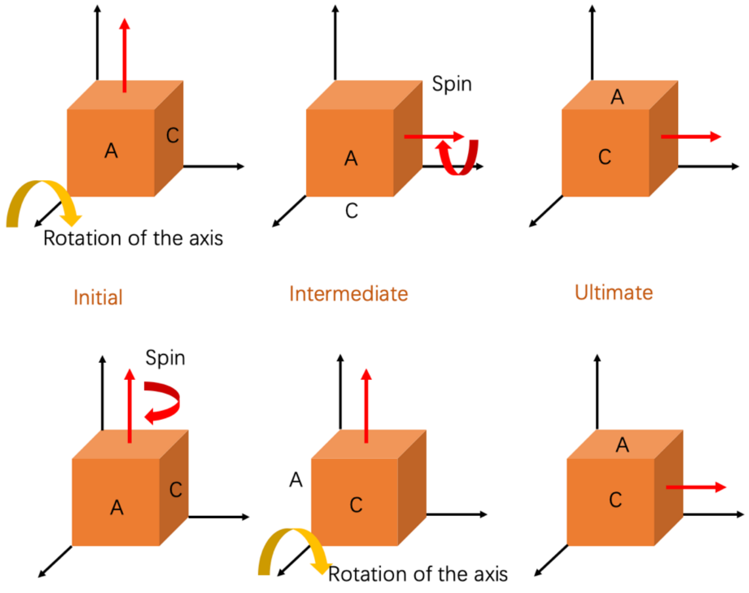

**FIG S1**. A cube undergoing different operations with different orders. Two orders of operations (top and bottom) give the same ultimate state.

The 3-dimentional rotation can be decomposed into 2 independent operations, axial rotation and spinning about its axis. Other choices of the rotational operations can affect the target state, thus to be order-dependent in general. For example, a cube undergoing the above two axial rotation and the spinning operations with different orders result in the same final state, i.e., the rotational operations are commutable,

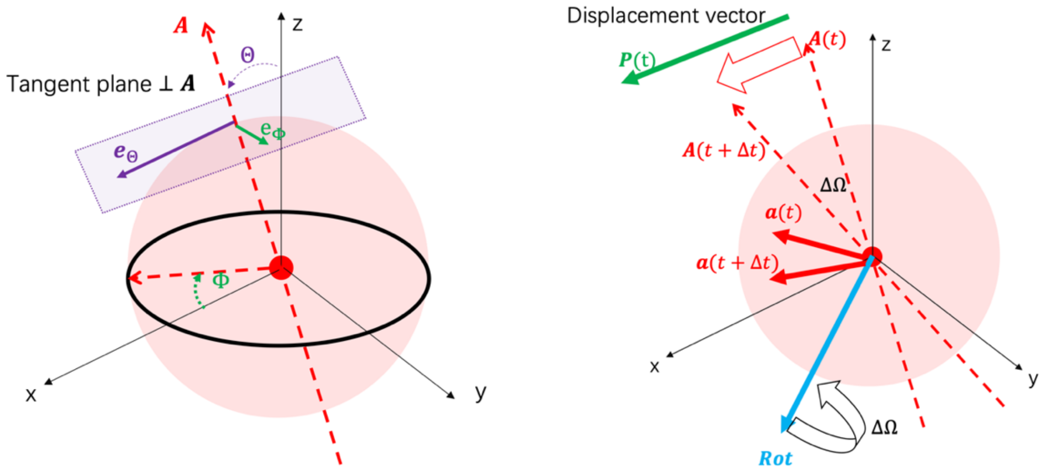


$$Rot_{Axis}\left(-\frac{\pi}{2}\right)\cdot Rot_{spin}\left(-\frac{\pi}{2}\right) = Rot_{spin}\left(-\frac{\pi}{2}\right)\cdot Rot_{Axis}\left(-\frac{\pi}{2}\right) \quad (S3)$$

**FIG S2.** (A)Spatial rotation of the axial vector of the spherical protein, $\boldsymbol{A}$, described by polar angle $\Theta$ (relative to the z-axis) and azimuthal angle $\Phi$ (relative to the x-axis). The tangent plane of the sphere perpendicular to $\boldsymbol{A}$ is uniquely determined, which can be spanned by $\boldsymbol{e}_\Theta$ and $\boldsymbol{e}_\Phi$.(B) Rotation of the axis $\boldsymbol{A}$ without spinning. The projection of the forces and the fluctuations together onto the tangent plane cause the rotation of the axis $\boldsymbol{A}$. The displacement unit-vector of $\boldsymbol{A}$, $\boldsymbol{P}$ and $\boldsymbol{A}$ define the rotational axis $\boldsymbol{Rot}$. An arbitrary vector $\boldsymbol{a}$ rotates about $\boldsymbol{Rot}$ by angle $\Delta\Omega$.

Angular displacements follow the Langevin dynamics

$$\Delta\Theta = -\frac{dt}{\zeta_{rot}}\sum_i \left((a_y F_{i,z} + a_z F_{i,y})e_{\Theta,x} + (a_z F_{i,x} + a_x F_{i,z})e_{\Theta,y} + (a_x F_{i,y} + a_y F_{i,x})e_{\Theta,z}\right)\frac{\mathbf{1}}{|\boldsymbol{a}|^2} + \sqrt{\frac{2k_B T dt}{\zeta_{rot}}}\,w/|\boldsymbol{a}| \quad (\boldsymbol{S4})$$

$$\Delta\Phi = \frac{dt}{\zeta_{rot}}\sum_i \left((a_y F_{i,z} + a_z F_{i,y})e_{\Phi,x} + (a_z F_{i,x} + a_x F_{i,z})e_{\Phi,y} + (a_x F_{i,y} + a_y F_{i,x})e_{\Phi,z}\right)\frac{\mathbf{1}}{|\boldsymbol{a}|^2} + \sqrt{\frac{2k_B T dt}{\zeta_{rot}}}\,w/|\boldsymbol{a}| \quad (\boldsymbol{S5})$$

The displacement vector is defined by

$$P_x = (\Delta\Theta e_{\Theta,x} + \Delta\Phi e_{\Phi,x})/\sqrt{\Delta\Theta^2 + \Delta\Phi^2}$$

$$P_y = (\Delta\Theta e_{\Theta,y} + \Delta\Phi e_{\Phi,y})/\sqrt{\Delta\Theta^2 + \Delta\Phi^2}$$

$$P_z = (\Delta\Theta e_{\Theta,z} + \Delta\Phi e_{\Phi,z})/\sqrt{\Delta\Theta^2 + \Delta\Phi^2}$$

Accordingly, the rotational axis is defined as

$$Rot_x = A_y P_z - A_z P_y$$
$$Rot_y = A_z P_x - A_x P_z$$
$$Rot_z = A_x P_y - A_y P_x$$

The resultant angle $\Delta\Omega = \sqrt{\Delta\Theta^2 + \Delta\Phi^2}$, an arbitrary vector $\boldsymbol{a}$ rotates about $\boldsymbol{Rot}$

$$\boldsymbol{a}(t + \Delta t) = \begin{pmatrix} \cos[\Delta\Omega] & -\sin[\Delta\Omega] & 0 \\ \sin[\Delta\Omega] & \cos[\Delta\Omega] & 0 \\ 0 & 0 & 1 \end{pmatrix} \begin{pmatrix} \boldsymbol{a}\cdot\boldsymbol{A} \\ \boldsymbol{a}\cdot\boldsymbol{P} \\ \boldsymbol{a}\cdot\boldsymbol{Rot} \end{pmatrix} \qquad (\boldsymbol{S6})$$

Specifically, the components of vector $\boldsymbol{a}$ are

$$a_x(t + \Delta t) = [\cos[\Delta\Omega]\,(\boldsymbol{a}_i \cdot \boldsymbol{A}) - \sin[\Delta\Omega]\,(\boldsymbol{a}_i \cdot \boldsymbol{P})]A_x + [\sin[\Delta\Omega]\,(\boldsymbol{a}_i \cdot \boldsymbol{A}) + \cos[\Delta\Omega]\,(\boldsymbol{a}_i \cdot \boldsymbol{P})]P_x + \boldsymbol{a}_i \cdot \boldsymbol{Rot}\, Rot_x$$

$$a_y(t + \Delta t) = [\cos[\Delta\Omega]\,(\boldsymbol{a}_i \cdot \boldsymbol{A}) - \sin[\Delta\Omega]\,(\boldsymbol{a}_i \cdot \boldsymbol{P})]A_y + [\sin[\Delta\Omega]\,(\boldsymbol{a}_i \cdot \boldsymbol{A}) + \cos[\Delta\Omega]\,(\boldsymbol{a}_i \cdot \boldsymbol{P})]P_y + \boldsymbol{a}_i \cdot \boldsymbol{Rot}\, Rot_y$$

$$a_z(t + \Delta t) = [\cos[\Delta\Omega]\,(\boldsymbol{a}_i \cdot \boldsymbol{A}) - \sin[\Delta\Omega]\,(\boldsymbol{a}_i \cdot \boldsymbol{P})]A_z + [\sin[\Delta\Omega]\,(\boldsymbol{a}_i \cdot \boldsymbol{A}) + \cos[\Delta\Omega]\,(\boldsymbol{a}_i \cdot \boldsymbol{P})]P_z + \boldsymbol{a}_i \cdot \boldsymbol{Rot}\, Rot_z$$

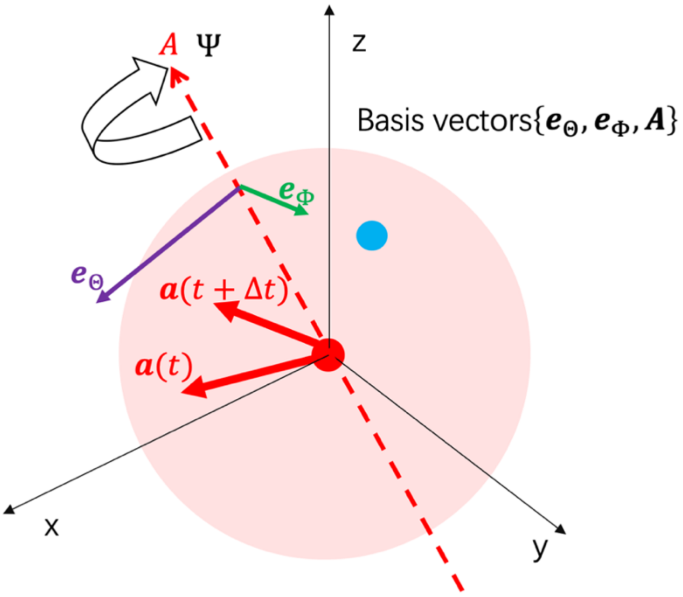


**FIG S3**. Spinning about the axis $\boldsymbol{A}$. The resulting spinning operation is conducted to the basis vectors $\{\boldsymbol{e}_\Theta, \boldsymbol{e}_\Phi, \boldsymbol{A}\}$. Vector $\boldsymbol{a}$ rotates about $\boldsymbol{A}$ by angle $\Psi$.

Finally, spinning of the sphere about its axis $\boldsymbol{A}$ follows

$$\Psi = \frac{dt}{\zeta_{rot}} \sum_i \left( (a_y F_{i,z} + a_z F_{i,y}) e_{\Psi,x} + (a_z F_{i,x} + a_x F_{i,z}) e_{\Psi,y} + (a_x F_{i,y} + a_y F_{i,x}) e_{\Psi,z} \right) \frac{\mathbf{1}}{|\boldsymbol{a}|^2}$$

$$+ \sqrt{\frac{2k_B T dt}{\zeta_{rot}}} w / \, |\boldsymbol{a}| \qquad (\boldsymbol{S7})$$

An arbitrary vector $\boldsymbol{a}$ rotates about $\boldsymbol{A}$

$$\boldsymbol{a}(t+\Delta t) = \begin{pmatrix} \cos[\Psi] & -\sin[\Psi] & 0 \\ \sin[\Psi] & \cos[\Psi] & 0 \\ 0 & 0 & 1 \end{pmatrix} \begin{pmatrix} \boldsymbol{a} \cdot \boldsymbol{e}_\Theta \\ \boldsymbol{a} \cdot \boldsymbol{e}_\Theta \\ \boldsymbol{a} \cdot \boldsymbol{A} \end{pmatrix}$$

The components of vector $\boldsymbol{a}$ are

$$a_x(t+\Delta t) = [\cos[\Psi]\,(\boldsymbol{a}_i \cdot \boldsymbol{e}_\Theta) - \sin[\Psi]\,(\boldsymbol{a}_i \cdot \boldsymbol{e}_\Phi)]\boldsymbol{e}_{\Theta,x} + [\sin[\Psi]\,(\boldsymbol{a}_i \cdot \boldsymbol{e}_\Theta) + \cos[\Psi]\,(\boldsymbol{a}_i \cdot \boldsymbol{e}_\Phi)]\boldsymbol{e}_{\Phi,x} + (\boldsymbol{a}_i \cdot \boldsymbol{A})\,A_x$$

$$a_y(t+\Delta t) = [\cos[\Psi]\,(\boldsymbol{a}_i \cdot \boldsymbol{e}_\Theta) - \sin[\Psi]\,(\boldsymbol{a}_i \cdot \boldsymbol{e}_\Phi)]\boldsymbol{e}_{\Theta,y} + [\sin[\Psi]\,(\boldsymbol{a}_i \cdot \boldsymbol{e}_\Theta) + \cos[\Psi]\,(\boldsymbol{a}_i \cdot \boldsymbol{e}_\Phi)]\boldsymbol{e}_{\Phi,y} + (\boldsymbol{a}_i \cdot \boldsymbol{A})\,A_y$$

$$a_z(t+\Delta t) = [\cos[\Psi]\,(\boldsymbol{a}_i \cdot \boldsymbol{e}_\Theta) - \sin[\Psi]\,(\boldsymbol{a}_i \cdot \boldsymbol{e}_\Phi)]\boldsymbol{e}_{\Theta,z} + [\sin[\Psi]\,(\boldsymbol{a}_i \cdot \boldsymbol{e}_\Theta) + \cos[\Psi]\,(\boldsymbol{a}_i \cdot \boldsymbol{e}_\Phi)]\boldsymbol{e}_{\Phi,y} + (\boldsymbol{a}_i \cdot \boldsymbol{A})\,A_z$$

The above dynamics can be briefly summarized as the following:

$$\dot{\boldsymbol{R}}_{COM} = -\frac{1}{\zeta} \nabla_{R_{COM}} H + \sqrt{\frac{2k_B T}{\zeta}} \boldsymbol{w} \qquad (\boldsymbol{S8})$$

$$\boldsymbol{A} \times \dot{\boldsymbol{\Omega}}_A |a| = -\frac{1}{\zeta_{rot}} \sum_i (a \times \boldsymbol{F}_i) \times \frac{\boldsymbol{A}}{|\boldsymbol{a}|} + \sqrt{\frac{2k_B T}{\zeta_{rot}}} (\boldsymbol{A} \times \boldsymbol{e}_{\boldsymbol{\Theta}} w_1 + \boldsymbol{A} \times \boldsymbol{e}_{\boldsymbol{\Phi}} w_2) \qquad (S9)$$

$$\boldsymbol{A} \cdot \dot{\boldsymbol{\Psi}} |a| = -\frac{1}{\zeta_{rot}} \sum_i (a \times \boldsymbol{F}_i) \cdot \frac{\boldsymbol{A}}{|\boldsymbol{a}|} + \sqrt{\frac{2k_B T}{\zeta_{rot}}} (w_3) \qquad (S10)$$

**C. Testing of the model**

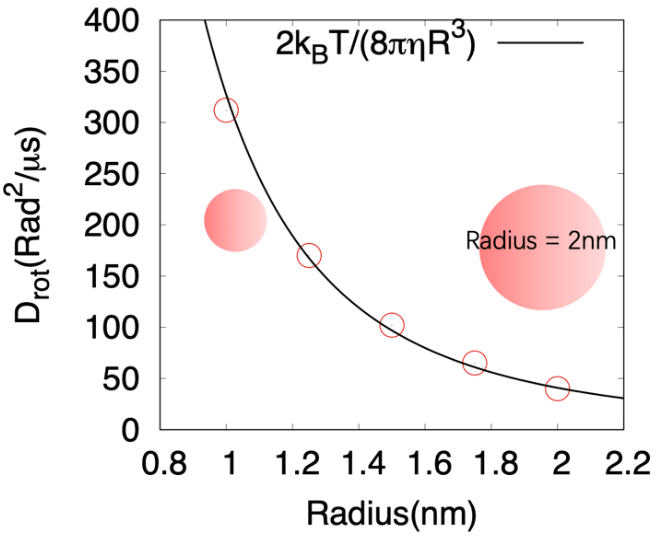


**FIG S4**. Rotation of the spherical proteins about a given axis. Based on the Einstein relation, the rotational diffusional coefficient is a function of the radius (black curve).

### 2. Restrained angular degree in diffusion

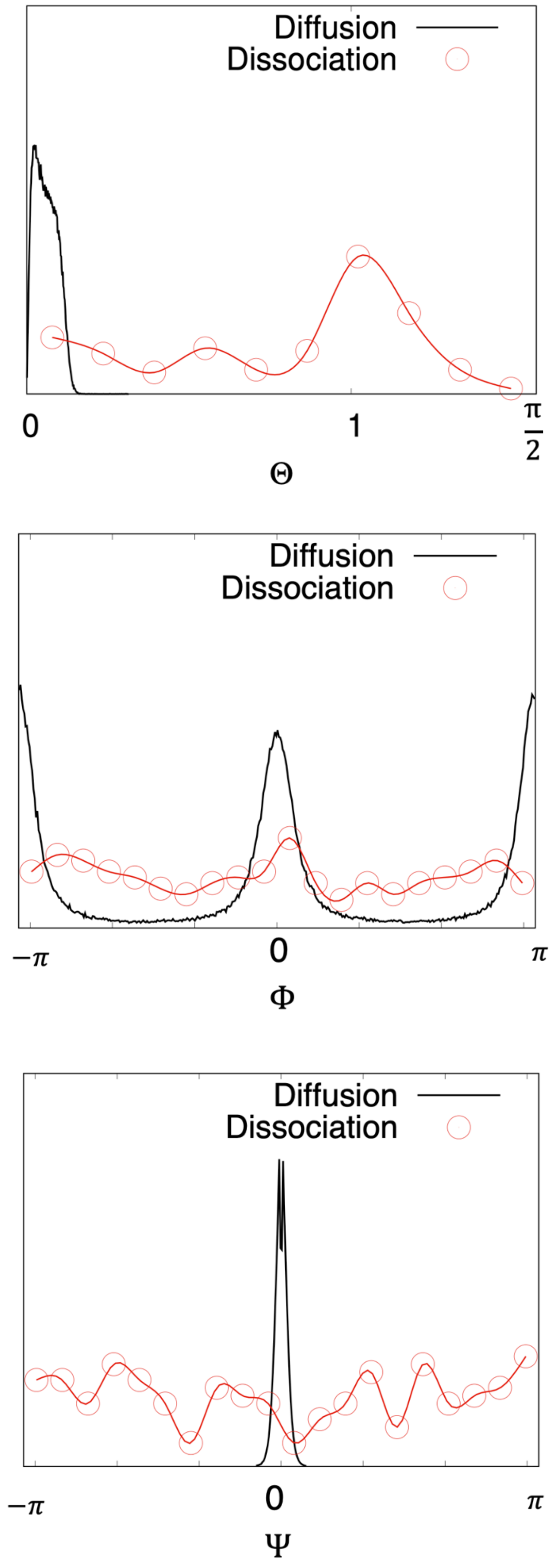

**FIG S5** Distributions of the rotational degrees of the spherical protein during diffusion obtained from the Langevin dynamics simulations. Spontaneous dissociation is led by the broken of the HB bonds, which requires much larger variation in $\Theta$ and $\Psi$. Diffusion along DNA requires the restrained angular degree to stabilize the protein-DNA HB contacts. For diffusion, the integral upper bound of $\Theta$ should take $0.1\pi$, the lower and upper bound of $\Psi$ take $-0.1\pi$ and $0.1\pi$.

### 3. Sequence-dependent stepping statistics along a random DNA sequence

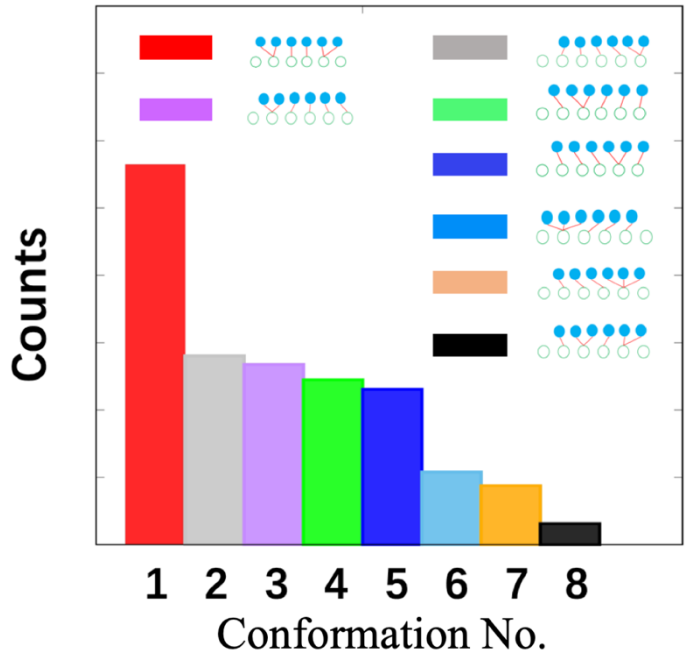


**FIG S6**. Typical HB configurations for protein sliding on a random sequence of length at 3000bp. The most probable configuration (red bar) is the same as that for protein stepping along the poly A sequence.

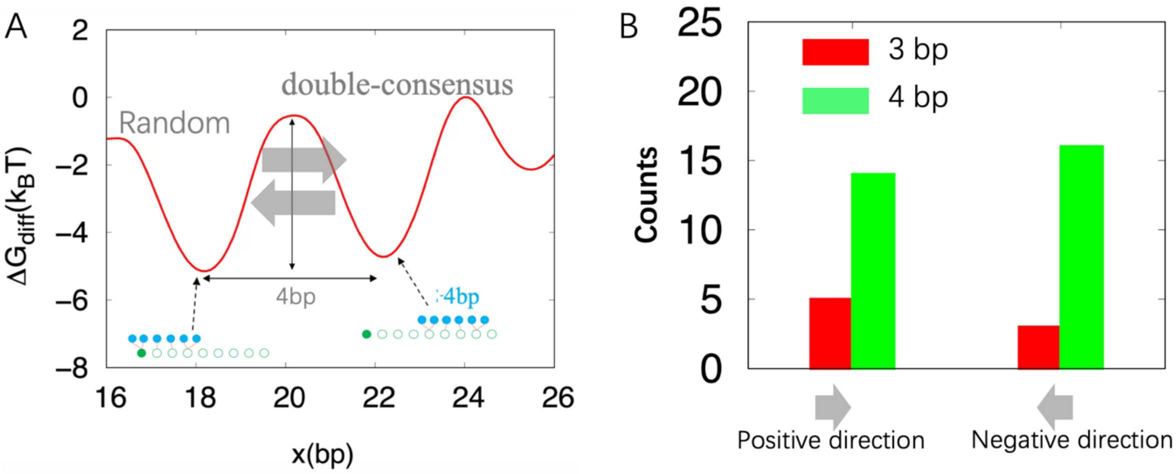


**FIG S7.** Comparing the HB positioning stepping size and separation between free energy minima. (A) Example of a short random sequence with double consensus sites (4bp in separation). The positive and negative stepping (grey arrows) from the 1st consensus site (at 18bp) to the 2nd one (at 22bp) accompany with the overcoming of the energy barrier ($\approx 5k_BT$). (B) HB positioning stepping sizes between the neighboring minima. The predominant HB stepping size (4bp) for both directions are the same as the separation between the neighboring minima. Therefore, the stepping size can be defined by the separation between the neighboring minima, which is experimentally measurable.

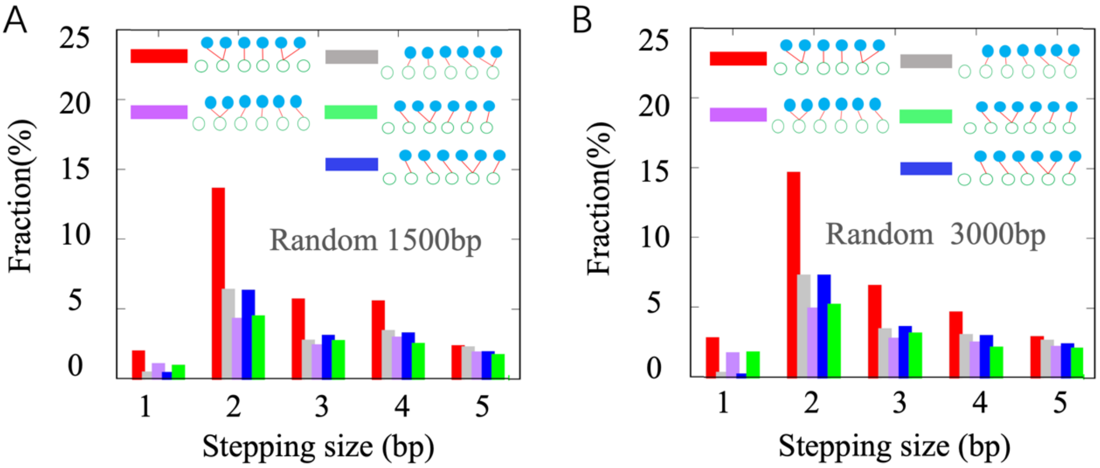


**FIG S8**. Convergence of the distributions of the stepping size. (A) Histogram for the fractions of the populations of the stepping sizes per cycle along random sequence at 1500 bp. (B) Histogram for the fractions of the populations of the stepping sizes sampled from the longer random sequence (3000 bp).

### 4. Multi-energy barriers give a rise to power law behavior

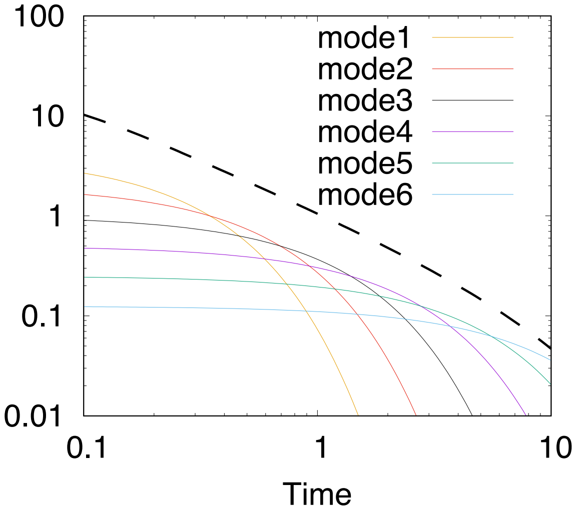


**FIG S9** The superposition of the multiple modes provides the power-law behavior in the stepping time. Each mode $ke^{-kt}$ corresponds to an energy barrier, e.g, the modes from 1 to 6 corresponding to rate k=4,2,1,0.5,0.25,0.125. The dashed line is the sum of 30 modes subjected to exponential distribution, which is approaching to power law as the number of the modes increases.

For a given energetic barrier $\Delta h$, the stepping time obeys exponential distribution $ke^{-kt}$, where the rate $k = k_0 e^{-\beta\Delta h}$. Thus, the continuum in energetic barriers $\Delta h$ leads to the superposition of various time distributions. Each exponential mode corresponds to an energy barrier $\Delta h$. We assume that the energetic barrier $\Delta h$ obeys the exponential distribution $f(\Delta h) = Ae^{-\Delta h/\mathcal{E}}$, where $\mathcal{E}$ is an energetic coefficient. The probability density of k can be obtained from $g(k) = f(\Delta h(k))\frac{\partial\Delta h}{\partial k}$, where $\Delta h = -\frac{1}{\beta}ln\frac{k}{k_0}$. Then the probability density of k is

$$g(k) = \frac{1}{\beta k}Ae^{\frac{1}{\beta\mathcal{E}}ln\frac{k}{k_0}} = Ck^{\frac{1}{\beta\mathcal{E}}-1}$$

The time distribution is the superposition of the continuous mode of $ke^{-kt}$,

$$\int_0^\infty g(k)ke^{-kt}dk = \int_0^\infty k^{\frac{1}{\beta\mathcal{E}}}e^{-kt}dk = t^{-(\frac{1}{\beta\mathcal{E}}+1)}\int_0^\infty u^{\frac{1}{\beta\mathcal{E}}+1-1}e^{-u}\,du = t^{-(\frac{1}{\beta\mathcal{E}}+1)}\Gamma(\frac{1}{\beta\mathcal{E}}+1)$$

where $\Gamma(\frac{1}{\beta\mathcal{E}}+1)$ is the Gamma function. The continuum in energetic barriers $\Delta h$ leads to power law

$$\rho(t) \propto \frac{1}{t^{\nu+1}} \qquad (S11)$$

where $\nu = \frac{1}{\beta\mathcal{E}}$.

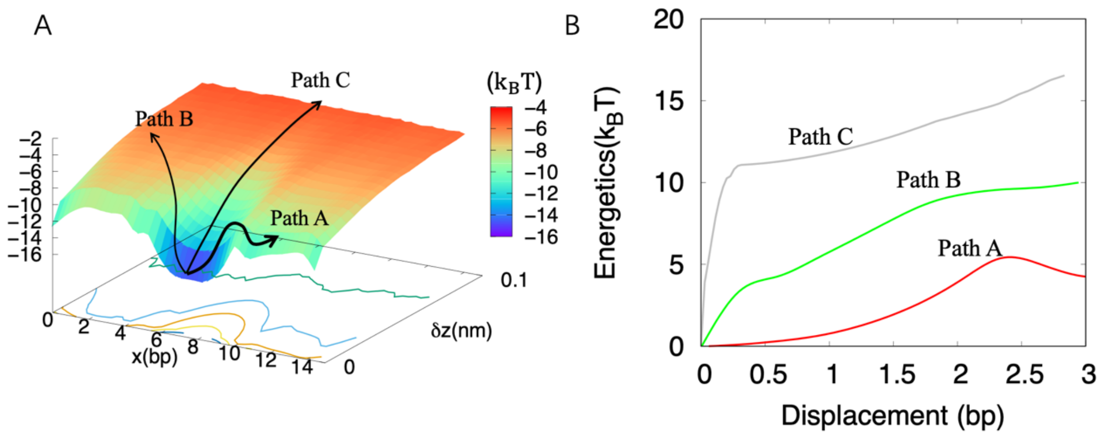


**FIG S10** (A)The dominant path A (via diffusion to flanking sites), the straightforward path B and the 'diffusion-coupled' path C for the protein dissociations.(B) Energy barriers for protein dissociations along the three paths.

### 5. Dissociation time versus DNA-length

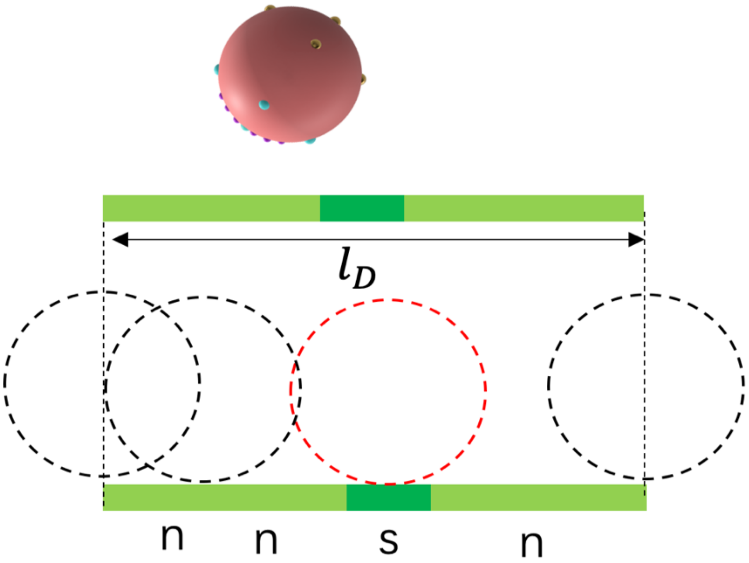


**FIG S11.** The width of the specific core center of the DNA is 1 bp, although the core covers about 6 base pairs. Then the non-specific sites center at $l_D - 1$ discrete bins (bp).

The population of the protein bound to DNA with HB contacts is proportional to $exp(-\Delta G_{diff})$. The dissociation rate $k(x) \propto exp(\Delta G_b)$. The mean dissociation rate is then

$$\bar{k} \propto \frac{(l_D - 1)exp(-\Delta G_{diff,n})exp(\Delta\Delta G_{b,n}) + exp(-\Delta G_{diff,s})exp(\Delta\Delta G_{b,s})}{(l_D - 1)exp(-\Delta G_{diff,n}) + exp(-\Delta G_{diff,s})} \quad \text{(S13)}$$

considering $l_D - 1$ sites for non-specific binding and one site for specific binding, with respective probability distributions weighted by $exp(-\Delta G_{diff,n})$ & $exp(-\Delta G_{diff,s})$, and respective dissociation rates weighted by \ $exp(\Delta\Delta G_{b,n})$ & $exp(\Delta\Delta G_{b,s})$.

For non-specific sites, we set approximately $\Delta G_{diff,n} \approx \Delta\Delta G_{b,n}$. Accordingly,

$$\bar{k} \propto \frac{l_D - 1 + exp(\Delta\Delta G_{b,s} - \Delta G_{diff,s})}{l_D - 1 + exp\left(\Delta\Delta G_{diff,n} - \Delta G_{diff,s}\right)} \quad \text{(S14)}$$

We have shown $\Delta\Delta G_{b,s} - \Delta G_{diff,s} = 1.2k_BT$ in the main text and $\Delta\Delta G_{diff,n} - \Delta G_{diff,s} = 7.1\, k_BT$. Therefore, the denominator of Eq.S14 is dominated by the factor $exp\left(\Delta\Delta G_{diff,n} - \Delta G_{diff,s}\right)$. The mean dissociation time is thus

$$\bar{\tau} \propto \frac{1}{l_D - 1 + exp(1.2)} \quad \text{(S15)}$$

One can derive $\Delta\Delta G$, the free energy difference of the specific site and non-specific site via the ratio of the dissociation rates, i.e., $\Delta\Delta G = -k_BTln\frac{k_{off,s}}{k_{off,n}}$, where $k_{off,s} = \frac{1}{\tau_s}$, and $k_{off,n} = \frac{1}{\tau_n}$. The free energy difference is hence $\Delta\Delta G = \Delta E_{bf}^{n-s} = \mathbf{3.3}k_BT$ for $l_D = 15bp$. For the core site (6bp), the recognition width is actually 1-bp. The free energy difference between the specific site and a nonspecific site can be calculated based on Eq.S15, i.e., $\Delta\Delta G = \Delta E_{bf}^{n-s} = 5.4k_BT$. In summary, calculating the free energy difference between the specific and non-specific sites via the dissociation rates gives a notable under-estimation of the free energy differences.
